# Distinct Neurexin-Cerebellin Complexes Control AMPA- and NMDA-Receptor Responses in a Circuit-Dependent Manner

**DOI:** 10.1101/2022.03.24.485585

**Authors:** Jinye Dai, Kif Liakath-Ali, Samantha Golf, Thomas C. Südhof

## Abstract

At mature CA1→subiculum synapses, alternatively spliced SS4+ variants of neurexin-1 (Nrxn1^SS4+^) and neurexin-3 (Nrxn3^SS4+^) enhance NMDA- and suppress AMPA-receptors, respectively. Both Nrxn1^SS4+^ and Nrxn3^SS4+^ act by binding to secreted cerebellin-2 (Cbln2) that in turn activates postsynaptic GluD1, which is homologous to AMPA- and NMDA-receptors. Whether neurexin-Cbln2-GluD1 signaling complexes have additional functions in synapse formation besides regulating NMDA- and AMPA-receptors, and whether they perform similar roles at other synapses, remains unknown. Using constitutive *Cbln2* deletions, we here demonstrate that at CA1→subiculum synapses, *Cbln2* performs no additional developmental functions besides regulating AMPA- and NMDA-receptors. Moreover, we show that low-level expression of Cbln1, which is functionally redundant with Cbln2, does not compensate for a synapse-formation function of Cbln2 at CA1→subiculum synapses. In exploring the generality of these findings, we found that in prefrontal cortex, Nrxn1^SS4+^-Cbln2 signaling selectively regulates NMDA-receptors, whereas Nrxn3^SS4+^-Cbln2 signaling has no apparent role. In contrast, in the cerebellum Nrxn3^SS4+^-Cbln1 signaling regulates AMPA-receptors, whereas now Nrxn1^SS4+^-Cbln1 signaling has no manifest effect. Thus, Nrxn1^SS4+^- and Nrxn3^SS4+^-Cbln1/2 signaling complexes generally control NMDA- and AMPA-receptors in different synapses without regulating synapse formation, but these signaling complexes are differentially active in diverse neural circuits.

## INTRODUCTION

Synaptic organizers are cell-adhesion molecules that direct the formation of synapses and shape their properties (Siddiqui and Craig, 2011; Ribic and Biederer, 2019; Sanes and Zipursky, 2020; Südhof, 2021; Kim et al., 2021; Graham and Duan, 2021). Multiple candidate synaptic organizers were described, among which neurexins and their multifarious ligands are arguably the best studied (reviewed in Noborn and Sterky, 2021; Gomez et al., 2021; Südhof, 2017; Kasem et al., 2018). Neurexins are presynaptic adhesion molecules that are encoded in mice by the *Nrxn1*, *Nrxn2*, and *Nrxn3* genes, each of which directs synthesis of longer α-neurexins and shorter β-neurexins from separate promoters (Tabuchi and Südhof, 2002). In addition, the *Nrxn1* gene (but not the *Nrxn2* and *Nrxn3* genes) contain a third promoter for even shorter Nrxn1γ (Sterky et al., 2017). All neurexin transcripts are extensively alternatively spliced at multiple sites, resulting in thousands of neurexin isoforms whose expression is tightly regulated (Lukacsovich et al., 2019; Nguyen et al., 2016; Ullrich et al., 1995; Fuccillo et al., 2015). Among the sites of alternative splicing of neurexins, splice site 4 (SS4) is possibly the most important because it regulates the interactions of neurexins with many of their ligands, including that of cerebellins (reviewed in Südhof, 2017). SS4 exhibits two variants that contain (SS4+) or lack (SS4-) a 30 residue insert, with only SS4+ neurexins binding to cerebellins.

Mammals contain four cerebellin genes (*Cbln1*-*4* in mice) that encode secreted multimeric C1q-domain proteins. Cerebellins function as trans-synaptic adaptors by connecting presynaptic neurexins to postsynaptic receptors (reviewed in Yuzaki, 2018; Matsuda, 2017). *Cbln1*, *Cbln2*, and *Cbln4* are broadly expressed in brain, whereas *Cbln3* is specific for cerebellar granule cells and requires Cbln1 for secretion (Bao et al., 2006; Miura et al., 2006). Remarkably, *Cbln1*, *Cbln2*, and *Cbln4* are not uniformly expressed in all neurons, but synthesized in restricted subsets of neurons (Seigneur and Südhof, 2017). For example, cerebellar granule cells express high levels of Cbln1 but only modest levels of Cbln2, excitatory entorhinal cortex neurons express predominantly Cbln4, and neurons in the medial habenula (mHb) express either Cbln2 or Cbln4 (Seigneur and Südhof, 2017). Although all cerebellins bind to presynaptic neurexins, they interact with different postsynaptic receptors: Cbln1 and Cbln2 bind to GluD1 and GluD2 (Matsuda et al., 2010), whereas Cbln4 binds to neogenin-1 (Neo1) and DCC (Wei et al., 2012; Haddick et al., 2014; Zhong et al., 2017). By connecting presynaptic neurexins to postsynaptic GluDs or to Neo1/DCC, cerebellins are thought to mediate trans-synaptic signaling and to organize synapses, but their precise functions are incompletely understood.

In the cerebellum, deletion of *Cbln1* or of its receptor GluD2 (gene symbol *Grid2*) throughout development causes a partial loss of parallel-fiber synapse numbers, and completely abolishes long-term plasticity (Hirai et al., 2005; Uemura et al., 2007; Rong et al., 2012a). Parallel-fiber synapses develop initially normally, but are subsequently lost (Kashiwabuchi et al., 1995; Kurihara et al., 1997; Hirai et al., 2005; Takeuchi et al., 2005).

Analyses of genetic deletions of *Cbln1*, *Cbln2*, and *Cbln4* outside of the cerebellum revealed behavioral changes and abnormal synaptic transmission with little or no loss of synapses (Kusnoor et al., 2010; Rong et al., 2012a; Otsuka et al., 2016; Seigneur et al., 2018; Seigneur and Südhof, 2018; Seigneur et al., 2021), suggesting a role in shaping synapse properties. For example, constitutive *Cbln1/2* double and *Cbln1/2/4* triple KO mice displayed major behavioral impairments but no synapse loss in the hippocampus at 2 months of age, although synapse loss developed over the next 4 months (Seigneur and Südhof, 2018). Moreover, a modest loss of synapses was detected in the striatum and prefrontal cortex at 6 months of age (Seigneur and Südhof, 2018). Furthermore, the constitutive deletion of Cbln2 produced abnormal compulsive behaviors in mice that resulted from insufficient activation of serotonergic neurons in the dorsal raphe, and could be reversed by administration of serotonergic agonists (Seigneur et al., 2021). Similarly, conditional inactivation of Cbln2 expression in the mHb led to major behavioral alterations and a rapid decline in mHb→interpeduncular nucleus synaptic transmission, but produced synapse loss only after 3 months (Seigneur et al., 2018).

Viewed together, these studies suggest that in all brain regions, cerebellins are essential for maintaining a normal behavioral repertoire and that they are not involved in the initial formation of synapses, but that their deletion in multiple brain regions causes a secondary loss of a subset of synapses. Contrary to this conclusion, however, an RNAi-induced suppression of Cbln2 expression was found to suppress formation of excitatory synapse numbers in the CA1 region of the hippocampus (Tao et al., 2018), which is puzzling since little Cbln2 can be detected in the CA1 region, and since constitutive KO mice at the same age do not exhibit a synapse loss (Seigneur and Südhof, 2017 and 2018). Similarly, an RNAi-induced suppression of Cbln4 expression was shown in the medial prefrontal cortex (mPFC) to cause an inhibitory synapse loss via a GluD1-dependent mechanism (Fossati et al., 2019), which is also puzzling since Cbln4 does not bind to GluD1 (Zhong et al., 2017; Cheng et al., 2016). Moreover, it has been reported that overexpression of human Cbln2 in mouse prefrontal cortex increases the spine density, implying an increase in synapse formation (Shibata et al., 2021). However, this observation also raised questions because again in *Cbln2* KO mice of the same age, little synapse loss is detected in the cortex (Seigneur and Südhof, 2018), and even in the cerebellum of *Cbln1* KO mice, the observed synapse loss is not accompanied by an equivalent decrease in spine density (Hirai et al., 2005). Moreover, the properties and the function of synapses were not actually measured by Shibata et al. (2021), making interpretations difficult. Finally, a synthetic synaptic organizer protein composed of Cbln1 fused to neuronal pentraxin 1 was shown to induce synapse formation in vivo (Suzuki et al., 2020), but in this experiment the binding partners of the synthetic protein were unclear, especially since little is known about the function of neuronal pentraxin 1, and the nature of the synaptogenic activity remained unexplored.

We previously demonstrated that at CA1→subiculum synapses, presynaptic neurexin-1 containing an insert in SS4 (Nrxn1^SS4+^) dominantly enhanced NMDA-receptor (NMDAR) EPSCs, whereas presynaptic neurexin-3 containing an insert in SS4 (Nrxn3^SS4+^) dominantly suppressed AMPA-receptor (AMPAR) EPSCs (Aoto et al., 2013; Dai et al., 2019). More recently, we showed that Nrxn1^SS4+^ and Nrxn3^SS4+^ both act by binding to Cbln2 which in turn binds to GluD1, indicating that at CA1→subiculum synapses, Nrxn1/3^SS4+^-Cbln2-GluD1 complexes mediate trans-synaptic signaling that controls NMDARs and AMPARs (Dai et al., 2021). No changes in synapse density were detected as a function of any of these manipulations – in fact, the massive increase in AMPAR EPSCs induced by the *Cbln2* deletion suggested that if a change in synapses occurred, it should have been an increase, not a decrease (Dai et al., 2021).

These results characterized a trans-synaptic signaling pathway that organized one particular synapse (CA1→subiculum synapses), but only this synapse was studied and it was only examined after it had fully developed, raising a series of questions. Specifically, does Cbln2 have additional essential roles in development at CA1→subiculum synapses? Since low levels of Cbln1 are also present at these synapses, is it possible that Cbln1 compensates for such additional functions in mature synapses? Furthermore, does Cbln2 function identically at different subtypes of CA1→subiculum synapses, where the properties of synapses formed on regular- and burst-firing neurons are quite different (Wojtowicz et al., 2010; Wozny et al., 2008a and 2008b)? More broadly and possibly more importantly, does a signaling pathway similar to the Nrxn1/3^SS4+^-Cbln2-GluD1 pathway operate at other synapses in brain, or is this pathway specific to CA1→subiculum synapses? To address these questions, we here first examined the role of Cbln2 and Cbln1 in CA1→subiculum synapses, and probed their function in relation to upstream Nrxn1^SS4+^ and Nrxn3^SS4+^ signals. We then studied the potential role of Nrxn1/3^SS4+^-Cbln1/2 signaling in two other paradigmatic synapses, namely Layer 2/3 to Layer 5/6 excitatory connections in the mPFC and parallel-fiber synapses in the cerebellum, which we investigated because previous work demonstrated a role for cerebellins in these brain regions. Our data suggest that the Nrxn1/3^SS4+^-Cbln1/2 signaling pathway has no role in synapse formation but functions to shape the NMDAR- and AMPAR-content at multiple types of synapses, and that different types of synapses exhibit distinct facets of this signaling pathway, such that in the mPFC, only the Nrxn1^SS4+^-Cbln2 signaling mechanism is present, whereas in the cerebellum, only the Nrxn3^SS4+^-Cbln1 signaling pathway operates.

## RESULTS

### Constitutive deletion of Cbln2 suppresses NMDARs and enhances AMPARs both at regular- and at burst-firing subiculum neuron synapses

Our previous conclusion that presynaptic Nrxn1^SS4+^ and Nrxn3^SS4+^ regulate postsynaptic NMDARs and AMPARs, respectively, via binding to Cbln2 but that Nrxn1, Nrxn3, and Cbln2 are not required for synapse formation relied on conditional manipulations at mature CA1→subiculum synapses (Dai et al., 2021). In contrast to these results, studies in the cerebellum (Hirai et al., 2005; Ito-Ishida et al., 2008; Rong et al., 2012a; Yuzaki, 2011) and the prefrontal cortex (Shibata et al., 2021) suggested a function for Cbln1 and Cbln2, respectively, in synapse formation, raising the question whether we might have overlooked such a role with conditional deletions. Moreover, in our experiments we did not differentiate between CA1→subiculum synapses on regular- and on burst-firing neurons that exhibit distinct forms of long-term plasticity (Wozny et al., 2008b). To explore whether Cbln2 may have an earlier developmental role in addition to its regulation of AMPARs and NMDARs at mature CA1→subiculum synapses, we examined the effect of the constitutive deletion of Cbln2. To determine whether Cbln2 may have distinct functions at synapses on regular- and burst-firing neurons, moreover, we studied these synapses separately at the same time (Figure. 1).

**Figure 1:**
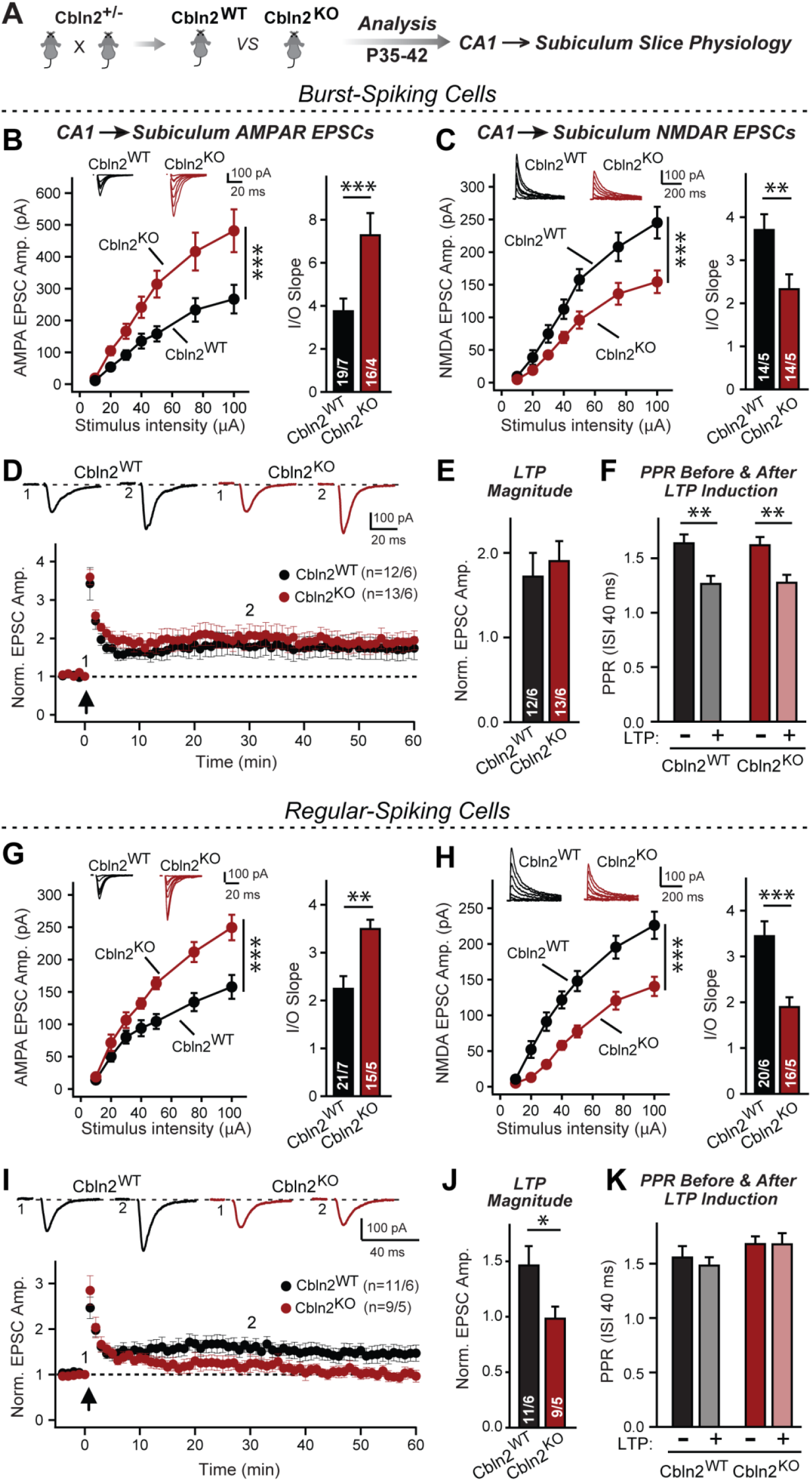
Constitutive *Cbln2* deletion increases AMPAR-EPSCs and suppresses NMDAR-EPSCs at CA1→subiculum synapses formed both on burst- and regular-spiking subiculum neurons, and blocks NMDAR-dependent LTP in regular-spiking neurons without affecting cAMP-dependent LTP in burst-spiking neurons. ***A*.** Experimental strategy for analysis of littermate wild-type and constitutive *Cbln2* KO mice. ***B*** *& **C***. Input/output measurements of evoked AMPAR- and NMDAR-EPSCs recorded from burst-spiking neurons in acute subiculum slices reveal that the *Cbln2* KO enhances AMPAR-EPSCs (B) but suppresses NMDAR-EPSCs (C). EPSCs were evoked by stimulation of CA1 axons in acute slices from littermate control and *Cbln2* KO mice at P35-42 (left, summary plots of input-output curves with sample traces on top; right, summary graph of input/output slopes). ***D****-**F***. The *Cbln2* KO had no effect on the presynaptic LTP typical for burst-spiking neurons that is induced by four 100 Hz/1 s stimulus trains with 10 s intervals under voltage-clamp (D, summary plot of AMPAR-EPSC amplitudes with sample traces on top; E, summary graph of the LTP magnitude (normalized EPSC amplitudes during the last 5 mins of recordings at least 30 min after LTP induction); F, summary graph of paired-pulse ratios before and after LTP induction as a measure of the release probability). ***G*** *& **H***. Same as B & C, but recorded from regular-spiking neurons. Note that the AMPAR-EPSC and NMDAR-EPSC phenotype of the *Cbln2* KO is identical in burst- and regular-spiking neurons. ***I****-**K***. The *Cbln2* KO abolishes NMDAR-dependent postsynaptic LTP that is typical for regular-firing subiculum neurons, and does not involve a change in PPR. Data are from experiments analogous to those described in D-F. All data are means ± SEM. Number of neurons/mice are indicated in bars. Statistical significance was assessed by unpaired two-tailed t-test or two-way ANOVA (*P≤0.05, **P≤0.01, and ***P≤0.001).

We generated and probed littermate WT and constitutive Cbln2 KO mice, and examined CA1→subiculum synaptic transmission in acute slices at postnatal day 35-42 (P35-42) (Figure 1A). In these experiments, we distinguished between regular- or burst-firing neurons in the subiculum by their electrical properties, stimulated axons emanating from the CA1 region, the major source of excitatory inputs into the subiculum (Bohm et al., 2018), and monitored EPSCs. In both regular- and burst-firing neurons, the constitutive Cbln2 deletion caused a large elevation (∼50%) in AMPAR-EPSC amplitudes and a similarly large decrease (∼50%) in NMDAR-EPSC amplitudes, as quantified in input/output curves to control for differences in stimulation efficiency (Figure 1B, 1C, 1G, 1H). These results exactly duplicated those obtained with conditional deletions, suggesting that the absence of Cbln2 throughout development did not produce an additional change in synaptic responses. Moreover, the finding that synapses on regular- and burst-firing neurons, the two different major types of excitatory synapses in the subiculum, are identically regulated by Cbln2 was confirmed in additional conditional deletion experiments (Figure S1).

Although CA1→subiculum synapses on regular- and burst-firing subiculum neurons are similar, they exhibit distinct forms of LTP, with the former expressing an NMDAR-dependent form of postsynaptic LTP, whereas the latter exhibits a presynaptic form of LTP (Wozny et al., 2008b). The Cbln2 deletion had no effect on presynaptic LTP in burst-firing neurons (Figure 1D, 1E) with a change in paired-pulse ratios (PPRs, Figure 1F) after induction, but abolished postsynaptic LTP in regular-firing neurons (Figure 1I, 1J) without a change of PPR after induction (Figure 1K). This deficit in LTP could be due to impaired LTP induction given the reduced NMDAR-response in Cbln2 KO mice, but we previously found that constitutive expression of Nrxn3^SS4+^ that regulates only AMPARs but not NMDARs also blocks LTP (Aoto et al., 2013), suggesting that the deficit in NMDAR-dependent LTP in Cbln2 KO mice could also be caused by a change in AMPAR trafficking. Consistent with the dramatic changes in AMPAR- and NMDAR-responses in Cbln2 KO mice, we observed a significant deficiency in contextual learning and memory in Cbln2 KO mice as monitored using the two-chamber avoidance test (Figure 2; see also Dai et al., 2019).

**Figure 2:**
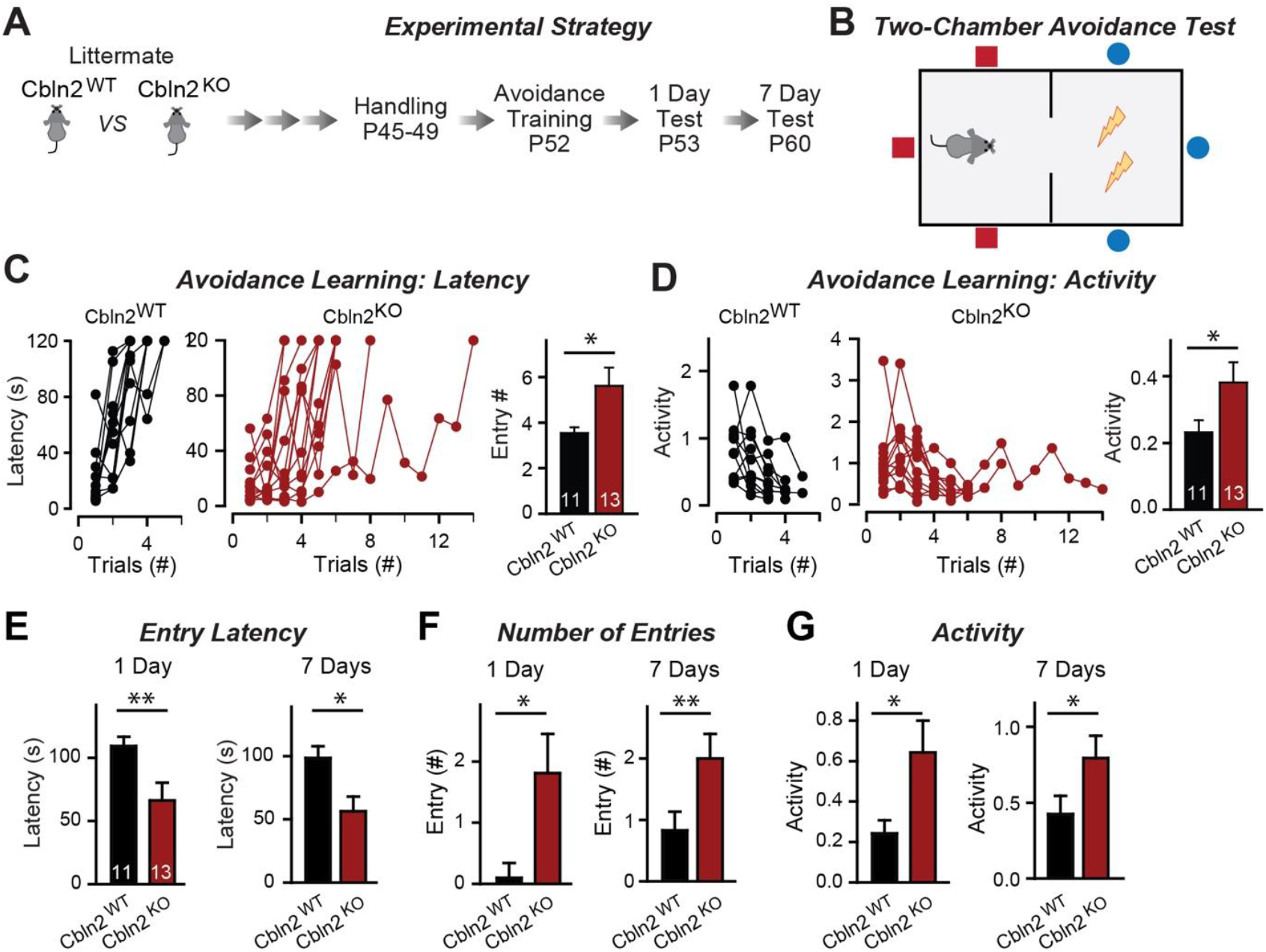
Constitutive *Cbln2* deletion impairs contextual memory in the two-chamber avoidance test. ***A*** & **B**. Experimental strategy of behavioral experiments utilizing littermate *Cbln2* KO and WT mice (A) and design of the two-chamber avoidance test in which mice receive mild electric foot shocks in the otherwise preferred darker chamber (B; Cimadevilla et al., 2001; Qiao et al., 2014). ***C*** *& **D***. *Cbln2* KO mice exhibit a delayed learning curve during two-chamber avoidance training. Mice learn to stay in the safe space by remembering visual cues to avoid the foot shock (C, trials for each mouse taking to learn when they remain in safe chamber for more than 2 mins (called latency; summary graphs shows number of entries); D, activity level in the safe chamber for each trial (summary graph shows activity level). ***E****-**G***. *Cbln2* KO severely decreases contextual memory in mice as measured by the two chamber avoidance test 1 day (left graphs) or 7 days (right graphs) after training (summary graphs of E, entry latencies; F, number of entries, and G, mouse activity). Data are means ± SEMs, the number of mice analyzed are depicted in the bars. Statistical analyses were performed by one-tail t-test (P ≤ 0.05; ** = P ≤ 0.01).

The finding that the constitutive and conditional deletion of Cbln2 produce the same synaptic phenotype suggests that the constitutive deletion, like the conditional deletion, does not produce differences in synapse formation, as would also be indicated by the dramatic increase in AMPAR-EPSC amplitudes induced by the Cbln2 deletion in both conditions. However, since cerebellins are broadly thought to mediate synapse formation (Kusnoor et al., 2010; Mishina et al., 2012; Matsuda, 2017; Seigneur and Sudhof, 2018; Yuzaki, 2018), we examined the overall synapse density in the subiculum as a function of the constitutive Cbln2 deletion using measurements of immunocytochemical staining intensity for vGluT1 and quantifications of synaptic protein levels as a proxy (Figure 3). The constitutive Cbln2 KO caused no change in vGluT1 staining intensity (Figure 3B, 3C) or in the levels of multiple synaptic proteins as assessed by quantitative immunoblotting (Figure 3D, 3E). Together with the lack of a decrease in AMPAR-mediated responses, these findings suggest that the constitutive Cbln2 KO ablating Cbln2 expression throughout development, similar to the conditional deletion in juvenile mice, does not decrease synapse numbers.

**Figure 3:**
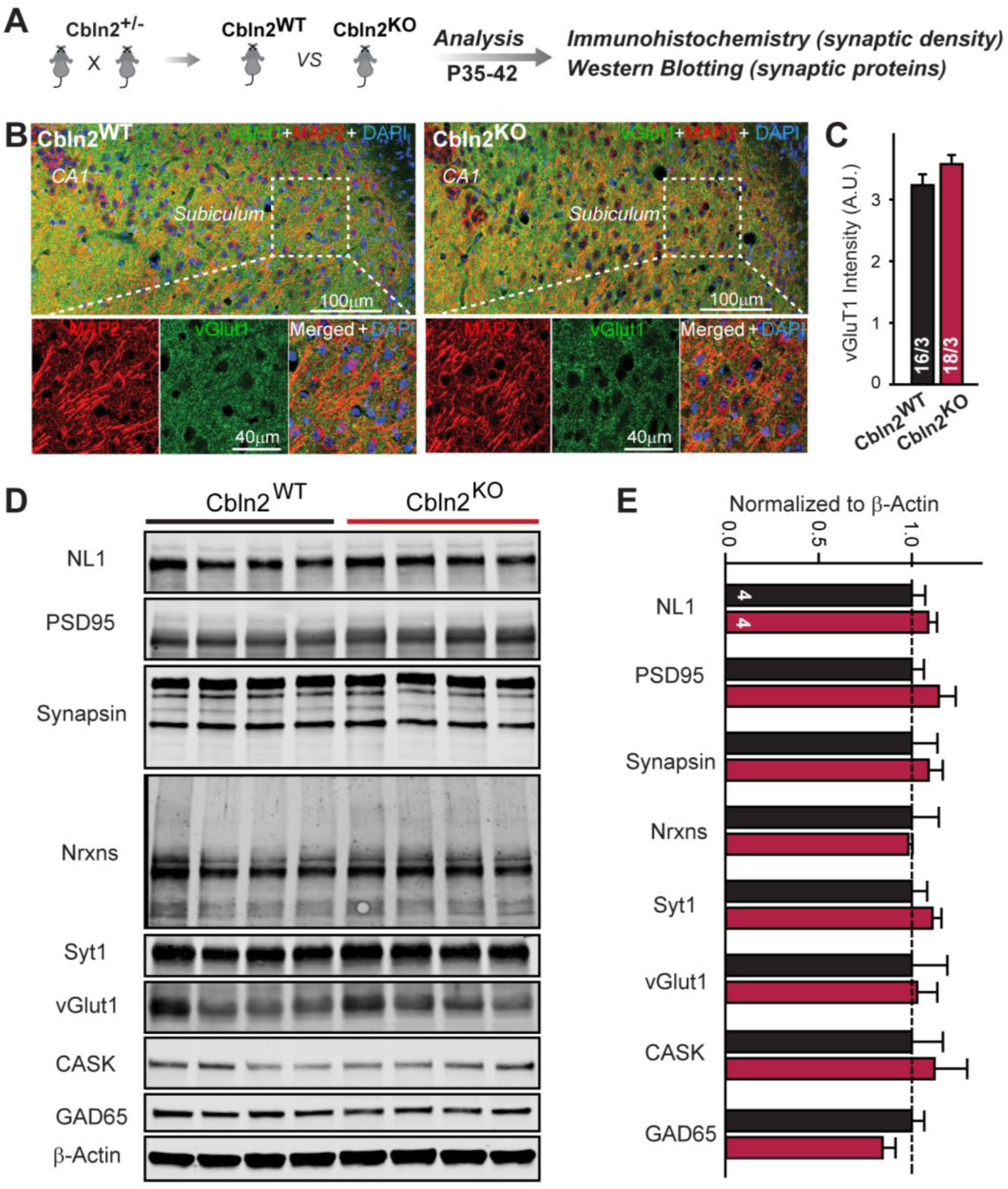
Constitutive *Cbln2* deletion does not alter the overall synapse density in the hippocampus. ***A.*** Experimental strategy for the analysis of littermate wild-type and constitutive *Cbln2* KO mice. ***B.*** Representative images of subiculum sections stained for vGluT1 as a proxy of synapse density, MAP2 as a proxy of neuronal density, and DAPI. **C.** The *Cbln2* KO does not cause a major loss of excitatory synapses in the subiculum as indicated by the vGluT1 staining intensity. ***D*** *& **E.*** The *Cbln2* KO also does not significantly alter the levels of synaptic proteins in the hippocampus. Protein levels were measured in hippocampal lysates by quantitative immunoblotting using fluorescent secondary antibodies (D, representative blots, please also see original full-sized immunoblots in Figure 3-source data 1; E, summary graph (levels are normalized for β-actin as an internal standard, and then to the controls to render results from multiple experiments comparable; n = 3 independent experiments). Data are means ± SEMs, the number of slices/mice or number of mice analyzed are depicted in the bars; statistical analyses by unpaired two-tailed t-test revealed no significant differences.

### Cbln2 regulates AMPARs and NMDARs via a trans-synaptic Nrxn1^SS4+^- and Nrxn3^SS4+^-dependent mechanism, respectively

We next set out to test whether the constitutive Cbln2 KO phenotype is due to the ablation of normally occurring presynaptic Nrxn1^SS4+^ and Nrxn3^SS4+^ signals, as suggested by previous studies (Aoto et al., 2013; Dai et al., 2019 and 2021). Quantifications of the alternative splicing of neurexins at SS4 in the CA1 region, subiculum, PFC, and cerebellum suggest that in the cerebellum, all neurexins are primarily expressed at SS4+ splice variants, whereas in the other three regions examined neurexins are expressed as a mixture of SS4+ and SS4-splice variants (Figure S2). Thus, a shift in alternative splicing of neurexins at SS4 could play a major regulatory role, as suggested previously (Ding et al., 2017; Fuccillo et al., 2015; Iijima et al., 2011). Therefore we used two experimental paradigms to induce such a shift and thereby to ask whether deletion of Cbln2 blocked the ability of Nrxn1^SS4+^ to enhance NMDAR-EPSCs and of Nrxn3^SS4+^ to suppress AMPAR-EPSCs.

First, we crossed constitutive Cbln2 KO mice with conditional Nrxn1^SS4+^ or Nrxn3^SS4+^ knockin mice (Aoto et al., 2013; Dai et al., 2019), and bilaterally infected the CA1 region of these double-mutant mice by stereotactic injections with AAVs encoding ΔCre (which retains the SS4+ splice variant) or Cre (which converts the presynaptic SS4+ splice variant into the SS4-variant) (Figure 4A). The Cbln2 deletion completely ablated the effect of the presynaptic Nrxn1^SS4+^ or Nrxn3^SS4+^ knockin on NMDAR- and AMPAR-ESPCs, respectively (Figure 4B, 4C). None of these manipulations altered PPRs, documenting that they did not influence the release probability (Figure 4D, 4E). These results confirm that Cbln2 is required for transduction of the presynaptic Nrxn1^SS4+^ or Nrxn3^SS4+^ signals into postsynaptic NMDAR and AMPAR responses, respectively.

**Figure 4:**
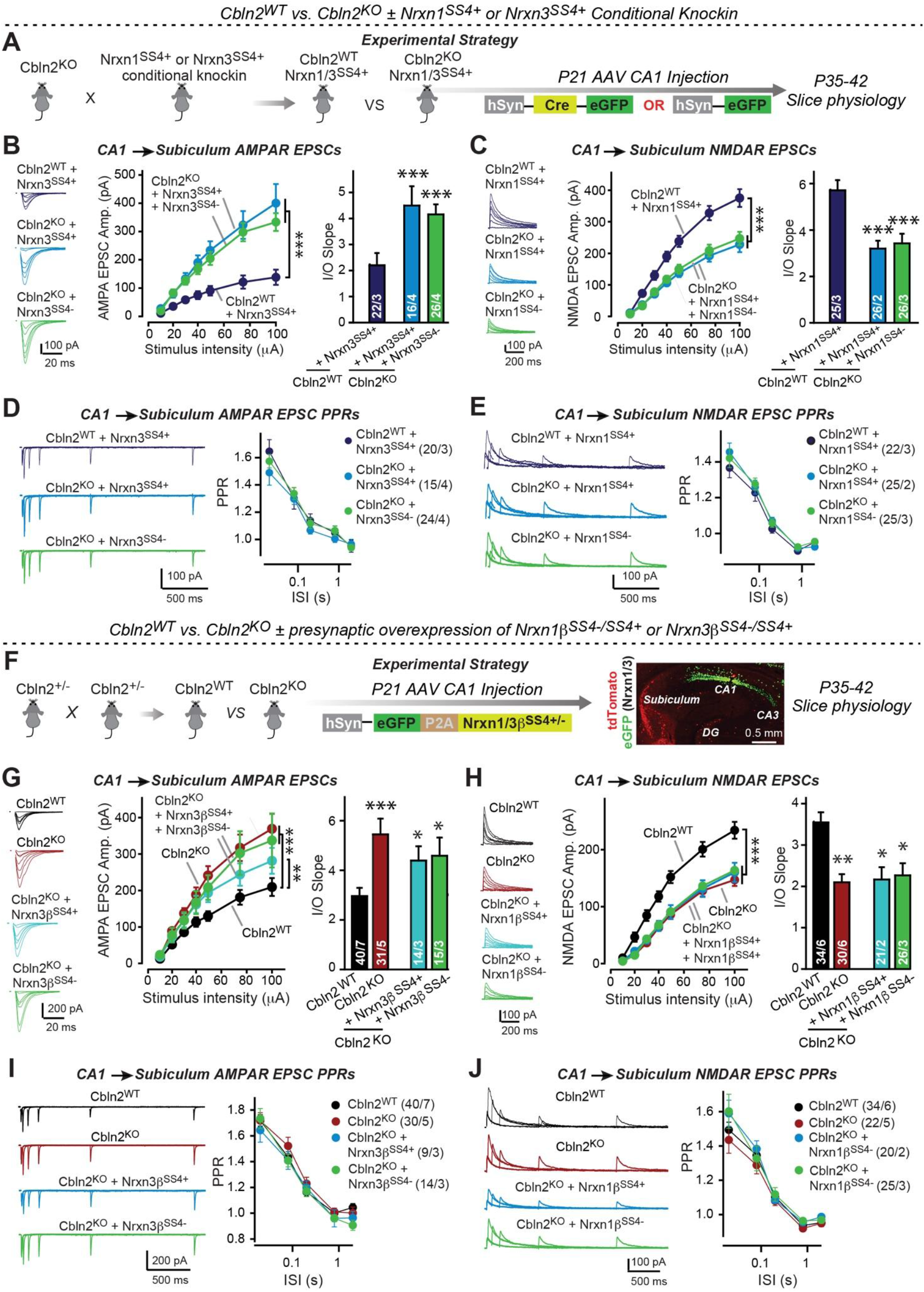
Constitutive *Cbln2* deletion occludes regulation of postsynaptic AMPAR- and NMDAR-EPSCs by presynaptic Nrxn1^SS4+^ and Nrxn3^SS4+^, respectively. ***A.*** Experimental strategy for testing whether the *Cbln2* deletion blocks the effects of Nrxn1^SS4+^ and Nrxn3^SS4+^ signaling. Constitutive *Cbln2* KO mice (Cbln2^KO^) were crossed with Nrxn1^SS4+^ and Nrxn3^SS4+^ knockin mice that constitutively express Nrxn1^SS4+^ and Nrxn3^SS4+^ splice variants, but that are converted into constitutively expressing Nrxn1^SS4-^ and Nrxn3^SS4-^ splice variants by Cre-recombinase (Dai et al., 2019). Three groups of mice were compared: 1. Cbln2^WT^ mice expressing Nrxn1^SS4+^ or Nrxn3^SS4+^, 2. Cbln2^KO^ mice expressing Nrxn1^SS4+^ and Nrxn3^SS4+^ in which presynaptic CA1 neurons were infected stereotactically at P21 with AAVs expressing inactive ΔCre (retains presynaptic Nrxn1^SS4+^ and Nrxn3^SS4+^ genotype); and 3. Nrxn1^SS4+^ and Nrxn3^SS4+^ in which presynaptic CA1 neurons were infected stereotactically at P21 with AAVs expressing active Cre (generates presynaptic Nrxn1^SS4-^ and Nrxn3^SS4-^ genotype). CA1→subiculum synapses were then analyzed in acute slices from these mice at P35-42. ***B.*** On the background of the *Cbln2* KO, knockin of Nrxn3^SS4+^ no longer suppresses AMPAR-ESPCs, nor does it reverse the increase in AMPAR-EPSCs induced by the Cbln2 KO at CA1→subiculum synapses (left, representative traces; middle, summary plot of the input/output relation; right, summary graph of the slope of the input/output relations). ***C.*** Similarly, Nrxn1^SS4+^ no longer enhances NMDAR-ESPCs on the background of the *Cbln2* KO, nor does it reverse the decrease in NMDAR-EPSCs induced by the Cbln2 KO (left, representative traces; middle, summary plot of the input/output relation; right, summary graph of the slope of the input/output relations). ***D*** *& **E**.* Constitutive expression of Nrxn1^SS4+^ and Nrxn3^SS4+^ alone or in combination with the *Cbln2* KO have no effect on the paired-pulse ratio of evoked AMPAR-EPSCs (D) or NMDAR-EPSCs (E) at CA1→subiculum synapses (left, sample traces; right, summary plots of PPRs). ***F.*** Alternative experimental strategy for testing whether the *Cbln2* deletion blocks the effects of Nrxn1^SS4+^ and Nrxn3^SS4+^ signaling. Analysing the epistatic relation of neurexin alternative splicing at SS4 with the *Cbln2* KO at CA1→subiculum synapses using viral overexpression of Nrxn1β^SS4+^ or Nrxn3β^SS4+^ in *Cbln2* KO mice. The CA1 region of constitutive *Cbln2* KO mice was bilaterally infected at P21 by stereotactic injections with AAVs expressing Nrxn1β^SS4+^, Nrxn1β^SS4-^, Nrxn3β^SS4+^, or Nrxn3β^SS4-^, and subiculum neurons were analyzed 2-3 weeks later. The representative image on the right depicts the signal for eGFP (which is co-expressed with the neurexins) in CA1 neurons after 2 weeks infection. ***G.*** On the background of the *Cbln2* KO, overexpression of Nrxn3β^SS4+^ again no longer suppresses AMPAR-ESPCs, nor does it reverse the increase in AMPAR-EPSCs induced by the *Cbln2* KO at CA1→subiculum synapses (left, representative traces; middle, summary plot of the input/output relation; right, summary graph of the slope of the input/output relations). ***H.*** Similarly, overexpressed Nrxn1β^SS4+^ no longer enhances NMDAR-ESPCs on the background of the *Cbln2* KO, nor does it reverse the decrease in NMDAR-EPSCs induced by the Cbln2 KO (left, representative traces; middle, summary plot of the input/output relation; right, summary graph of the slope of the input/output relations). ***I*** *& **J.*** Overexpression of any neurexin has no effect on the paired-pulse ratio of evoked AMPAR-EPSCs (I) or NMDAR-EPSCs (J) (left, sample traces; right, summary plots of PPRs). Data are means ± SEM. Number of neurons/mice are indicated in bars. Statistical significance was assessed by unpaired two-tailed t-test comparing to control and two-way ANOVA (*P≤0.05, **P≤0.01, and ***P≤0.001).

Second, we overexpressed Nrxn1β^SS4+^ or Nrxn3β^SS4+^ in the presynaptic CA1 region in constitutive Cbln2 KO mice *in vivo* using stereotactic bilateral injections of AAVs (Figure 4F). We previously showed that overexpression of Nrxn1β^SS4+^ in wild-type CA1 neurons increases NMDAR- but not AMPAR-EPSCs at CA1→subiculum synapses, whereas overexpression of Nrxn3β^SS4+^ in wild-type CA1 neurons suppresses AMPAR- but not NMDAR-EPSCs in the same synapses (Dai et al., 2019). When we tested the effect of Nrxn1β^SS4+^ or Nrxn3β^SS4+^ in constitutive Cbln2 KO mice, however, Nrxn1β^SS4+^ no longer increased NMDAR-EPSCs and Nrxn3β^SS4+^ no longer suppressed AMPAR-EPSCs (Figure 4G, 4H). None of these manipulations altered PPRs, demonstrating that they did not affect presynaptic properties (Figure 4I, 4J). Viewed together, these data suggest that Cbln2 transduces presynaptic Nrxn1^SS4+^ and Nrxn3^SS4+^ signals into distinct postsynaptic receptor responses at CA1→subiculum synapses.

### Double deletion of Cbln1 and Cbln2 produces the same phenotype as deletion of Cbln2 alone

Up to this point, our results indicate that Cbln2 functions both at regular- and at burst-firing neuron synapses in the subiculum to control AMPARs and NMDARs without being required for synapse formation. However, in these and earlier experiments we only studied Cbln2, but quantifications show that Cbln1 is also expressed in the subiculum, albeit at lower levels (Figure S3). Cbln1 and Cbln2 have biochemically nearly indistinguishable properties, suggesting that they are functionally redundant. The finding that Cbln1 is also expressed in the subiculum raises the possibility that the observed Cbln2 KO phenotype reflects only those functions of Cbln2 that are most sensitive to a decrease in overall Cbln1/2 levels, and that the remaining Cbln1 could occlude other phenotypes. To address this concern, we generated conditional Cbln1/2 double KO mice and analyzed the effect of the double Cbln1/2 deletion in the subiculum by electrophysiology, using a more expansive array of measurements to ensure that no effects were overlooked (Figure 5A).

**Figure 5:**
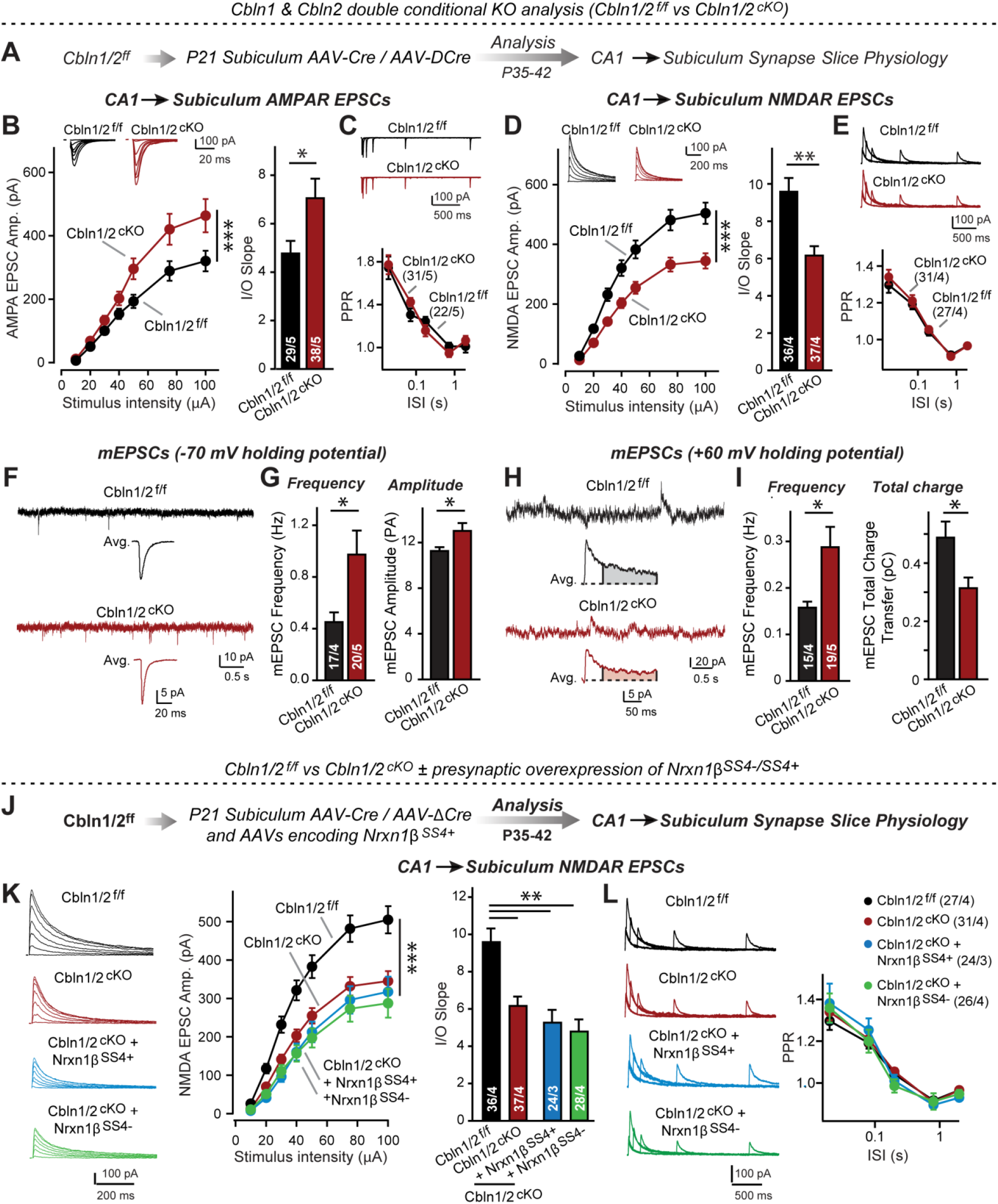
*Cbln1* and *Cbln2* double KO in the subiculum phenocopies the *Cbln2* single KO in CA1→subiculum synapses. ***A.*** Experimental strategy. AAVs encoding Cre or ΔCre (as a control) were stereotactically injected into the subiculum of conditional KO mice at P21, and mice were analyzed by slice physiology 2-3 weeks later. ***B****-**E***. Input/output measurements of evoked EPSCs recorded from combined burst- and regular-spiking neurons in acute subiculum slices reveal that the conditional Cbln2 KO enhances AMPAR-EPSCs (B) without changing the paired-pulse ratio of AMPAR-EPSCs (C) but suppresses NMDAR-EPSCs (D), again without changing the paired-pulse ratio of NMDAR-EPSCs (E). Sample traces are shown above the respective summary plots and graphs. ***F****-**I.*** Analyses of mEPSCs recorded at -70 mV and +60 mV holding potentials from burst- and regular-firing neurons in the subiculum after deletion of both Cbln1 and Cbln2 reveal an increase in mEPSC frequency measured at both holding potentials, but a decrease in charge transfer only of mEPSCs monitored at a +60 mV holding potential consistent with the decreased NMDAR-EPSC amplitude detected during input/output measurements (F, sample traces; G, bar graphs of the mEPSC frequency and amplitude, respectively; H & I, same as F & G but for recordings at +60 mV). ***J.*** Experimental strategy. The subiculum region of Cbln1/2^cKO^ was bilaterally infected at P21 by stereotactic injections of AAVs expressing ΔCre-eGFP (Cbln1/2^f/f^) or Cre-eGFP (Cbln1/2^cKO^), and then two weeks later cohorts of mice injected with Cre were further injected into the CA1 region with AAVs expressing Nrxn1β^SS4+^ or Nrxn1β^SS4-^. Mice were then analyzed at P49-P56 by acute slice electrophysiology. ***K.*** Overexpressed Nrxn1β^SS4+^ no longer enhances NMDAR-ESPCs on the background of the double Cbln1/2 cKO, nor does it reverse the decrease in NMDAR-EPSCs induced by the double Cbln1/2 cKO (left, representative traces; middle, summary plot of the input/output relation; right, summary graph of the slope of the input/output relations). ***L.*** Conditional deletion of both Cbln1 and Cbln2 without or with presynaptic overexpression of Nrxn1β^SS4+^ or Nrxn1β^SS4-^ does not alter paired-pulse ratios of NMDAR EPSCs. Left panels show sample traces; right panels summary plots of the paired-pulse ratio as a function of the interstimulus interval. Data are means ± SEMs; the number of cells/mice are depicted in the bars. Statistical analyses were performed by two-way ANOVA or unpaired two-tailed t-test comparing KOs to WT (*P ≤ 0.05; **P ≤ 0.01, and ***P ≤ 0.001).

Measurements of NMDAR-EPSCs and AMPAR-ESPCs elicited by stimulation of CA1 axons revealed the same phenotype in Cbln1/2 double conditional KO subiculum as the conditional and constitutive Cbln2-only deletion: A decrease in NMDAR-responses and an increase in AMPAR-responses (Figure 5B, 5D). These phenotypes were validated using input/output measurements to control for variabilities in the position of the stimulating electrode, and were due to a postsynaptic mechanism, as described before, since the PPRs did not change (Figure 5C, 5E). We also measured spontaneous mEPSCs as an indirect measure of synaptic activity and synapse numbers, and monitored mEPSCs at two holding potentials (−70 mV and +60 mV) to capture the contributions of both AMPARs and NMDARs to the mEPSCs (Figure 5F-5I). mEPSCs monitored at -70 mV are exclusively mediated by AMPARs, whereas mEPSCs monitored at +60 mV contain contributions of both AMPAR and NMDAR activation. At both holding potentials, the mEPSC frequency was massively enhanced (∼100-130% increase) by the Cbln1/2 double KO, presumably because of the increased AMPAR-responses leads to increased detection of mEPSCs at both holding potentials. Importantly, the average mEPSC amplitude was increased at the - 70 mV holding potential but the average mEPSC total charge transfer decreased at the +60 mV, consistent with the observation that the double Cbln1/2 KO increases AMPAR- but decreases NMDAR-responses (Figure 5B, 5D).

Finally, we asked whether the phenotype of the double Cbln1/2 KO might be more sensitive to manipulations of neurexins than that of the Cbln2 single KO. Focusing on Nrxn1 and NMDARs, we found that as with the single deletion of Cbln2, NMDAR EPSCs were no longer altered upon presynaptic overexpression of Nrxn1β containing or lacking an insert in SS4 (Figure 5J-5L). Overall, these data suggest that the Cbln1/2 double deletion has the same overall phenotype as the Cbln2 single deletion, with a dramatic change in AMPAR- and NMDAR-EPSCs but no apparent changes in presynaptic release probability.

### Nrxn1^SS4+^-Cbln2 complexes upregulate NMDARs in the prefrontal cortex, while Nrxn3^SS4+^-Cbln2 complexes have no effect

Our studies in two different CA1→subiculum synapses, described here and previously (Aoto et al., 2013 and 2015; Dai et al., 2019 and 2021), show that Nrxn1^SS4+^-Cbln2 complexes upregulate NMDARs whereas Nrxn3^SS4+^-Cbln2 complexes downregulate AMPARs. Does this trans-synaptic signaling pathway also operate in non-subiculum synapses, or is this a specific feature of subiculum synapses?

To address this question, we conditionally deleted Cbln2 from the mPFC (Figure 6A), which was chosen because it exhibits robust expression of Cbln2 (Figure S3) and because Cbln2 is particularly interesting in the mPFC due to its human-specific regulation of expression in this brain region (Shibata et al., 2021). We stereotactically injected AAVs encoding ΔCre or Cre into the mPFC of Cbln2 conditional KO mice at P21, and analyzed layer2/3 (L2/3) → layer5/6 (L5/6) synapses in acute slices at P35-42 (Figure 6A). For this purpose, we placed the stimulating electrode close to L2/3 neurons and recorded from L5/6 pyramidal neurons (Figure 6B).

**Figure 6:**
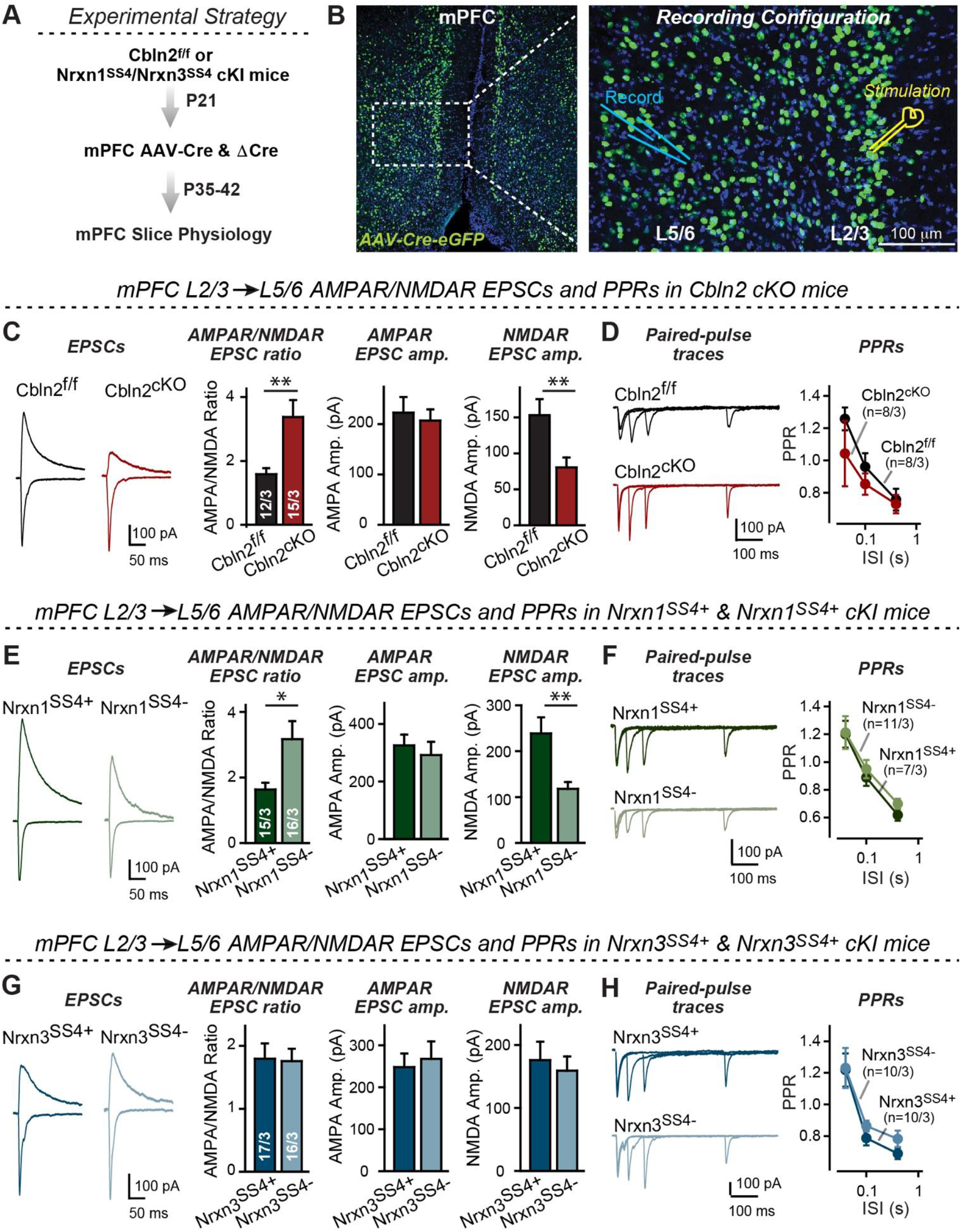
Nrxn1^SS4+^-Cbln2 signaling controls NMDAR-EPSCs but not AMPAR-EPSCs in the medial prefrontal cortex (mPFC), whereas Nrxn3^SS4+^-Cbln2 signaling does not regulate either AMPAR- or NMDAR-EPSCs in the mPFC. ***A & B***. Experimental strategy (left, flow diagram of the experiments; middle and right, Analysis strategies of Cbln2/Nrxn1-SS4/Nrxn3-SS4 conditional KO. Right, the mPFC region of Cbln2^cKO^ was bilaterally infected at P21 by stereotactic injections of AAVs expressing ΔCre-eGFP (Cbln2^f/f^) or Cre-eGFP (Cbln2^cKO^), and L5/6 pyramidal neurons in the prelimbic cortex (PL) region were analyzed 2-3 weeks later (A). The stimulation electrode was placed in L2/3 (B). ***C.*** Left, sample traces of evoked AMPAR- and NMDAR-EPSCs at Cbln2^f/f^ and Cbln2^cKO^ mPFC brain slices; Right, statistics of AMPA/NMDA ratios, AMPAR-EPSCs amplitude, and NMDAR-EPSCs amplitude. ***D.*** Left, sample traces of paired-pulse measurements from each condition; Right, summary plots of PPRs. ***E &F***. Same as C & D, but recorded from Nrxn1^SS4+^ knockin mice in which ΔCre retains a constitutive expression of Nrxn1-SS4+ splice variants, whereas Cre converts the Nrxn1-SS4+ variants into constitutive Nrxn1-SS4-variants. ***G & H***. Same as C & D, but recorded from Nrxn3^SS4+^ knockin mice in which ΔCre retains a constitutive expression of Nrxn3-SS4+ splice variants, whereas Cre converts the Nrxn3-SS4+ variants into constitutive Nrxn3-SS4-variants. Data are means ± SEM. Number of neurons/mice are indicated in bars. Statistical significance was assessed by unpaired two-tailed t-test or two-way ANOVA (*P≤0.05, **P≤0.01, and ***P≤0.001).

The conditional Cbln2 deletion produced a massive increase (∼100%) in the AMPAR/NMDAR ratio of L2/3→L5/6 synaptic transmission in the mPFC (Figure 6C). This increase was due to a large reduction (∼50%) in NMDAR-EPSC amplitudes without a change of AMPAR-EPSC amplitudes (Figure 6C). Again, we observed no changes in PPRs (Figure 6D).

These data suggest that, in L2/3→L5/6 synapses of the mPFC, Cbln2 surprisingly operates only as a regulator of NMDARs but not of AMPARs (Figure 6C, 6D). Is the function of Cbln2 in the mPFC also downstream of neurexins? To examine this question, we investigated the effect of the constitutive expression of Nrxn1^SS4+^ or Nrxn3^SS4+^ at L2/3→L5/6 synapses in the mPFC. We bilaterally infected the mPFC of Nrxn1^SS4+^ or Nrxn3^SS4+^ conditional knockin mice (Aoto et al., 2013; Dai et al., 2019) by stereotactic injections with AAVs encoding ΔCre (which retains the SS4+ variant) or Cre (which converts SS4+ variants into SS4-variants). Consistent with the Cbln2 KO results, only the presynaptic Nrxn1^SS4+^ knockin produced a phenotype, whereas the Nrxn3^SS4+^ knockin had no effect (Figure 6E-6H). Specifically, constitutive expression of Nrxn1^SS4+^ deletion produced a large increase (∼100%) in the AMPAR/NMDAR ratio due to a large decrease (∼100%) in the NMDAR-EPSC amplitudes but not AMPAR-EPSC, and this phenotype was abolished by conversion of Nrxn1^SS4+^ to Nrxn1^SS4-^ (Figure 6E). In contrast, the constitutive expression of Nrxn3^SS4+^ had no effect on the AMPAR/NMDAR ratio or either AMPAR-EPSC or NMDAR-EPSC amplitudes (Figure 6G). Again, none of these manipulations altered PPRs, documenting that they did not influence the release probability (Figure 6F, 6H). These results are consistent with the Cbln2 KO findings in the mPFC, validating the Nrxn1^SS4+^→Cbln2→NMDAR signaling pathway in the mPFC in the absence of the Nrxn3^SS4+^→Cbln2→AMPAR signaling pathway that we also observed in the subiculum.

### In the cerebellum, Nrxn3^SS4+^-Cbln1 complexes suppress AMPARs, whereas Nrxn1^SS4+^-Cbln1 complexes have no effect

Cerebellins were discovered in the cerebellum, with constitutive deletions of Cbln1 or of its receptor GluD2 causing a marked loss of parallel-fiber synapses (Hirai et al., 2005; Kashiwabuchi et al., 1995; Kurihara et al., 1997; Takeuchi et al., 2005). However, it is unclear whether this loss of synapses (that starts after synapses are initially formed) reflects a direct function of Cbln1 in synapse formation or represents an indirect effect of a change in AMPARs to which parallel-fiber synapses may be particularly sensitive (note that parallel-fiber synapses do not express functional NMDARs; Llano et al., 1991; Perkel et al., 1990). In the first case, Cbln1 would perform a function in the cerebellum that differs from that of Cbln2 in the subiculum; in the second case, Cbln1 would also regulate AMPARs in parallel-fiber synapses in a function that would be the same as the role we described for the subiculum, and that should become detectable in synapses after they have been formed.

To address this question, we stereotactically infected lobes 4-5 of the cerebellum of *Cbln1* conditional KO mice at P21 with AAVs encoding ΔCre or Cre, and analyzed parallel-fiber to Purkinje cell (PF-PC) synaptic transmission at P35-42 (Figure 7A). Strikingly, the *Cbln1* deletion increased the AMPAR-EPSC input/output curve and its slope (Figure 7C), without changing the coefficient of variation (CV), indicating that it did not influence the release probability (Figure 7D).

**Figure 7:**
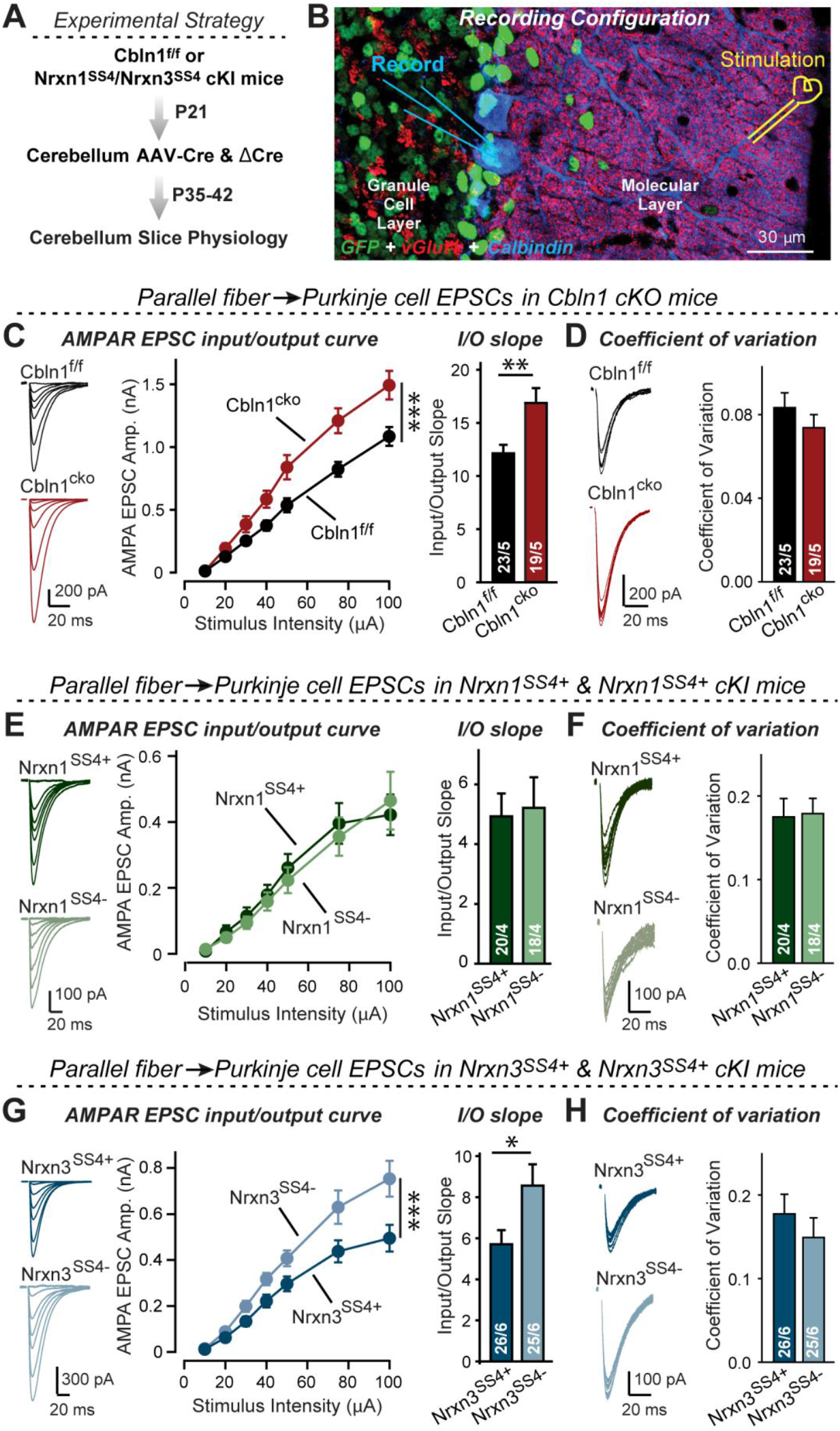
Nrxn3^SS4+^-Cbln1 signaling controls AMPAR-EPSCs in the cerebellum, but in this brain region Nrxn1^SS4+^-Cbln1 signaling has no effect. ***A.*** Experimental workflow for analyzing the effect of the *Cbln1* cKO or of the conditional Nrxn1^SS4+^ or Nrxn3^SS4+^ knockin on parallel-fiber synaptic transmission in the cerebellum. Note that the expression of ΔCre in Nrxn1^SS4+^ or Nrxn3^SS4+^ knockin mice retains the constitutive expression of their SS4+ splice variants, whereas the expression of Cre converts SS4+ into a constitutive SS4-splice variant. ***B.*** Image of a cerebellar cortex section (lobes 4-5) from *Cbln1* cKO mouse in which these lobes were infected at P21 by stereotactic injections of AAVs expressing ΔCre-eGFP (Cbln1^f/f^) or Cre-eGFP (Cbln1^cKO^). Sections were analyzed at P35 by slice physiology; the positions of the recording electrode in the patched Purkinje cells and of the stimulation electrode in the granule cell layer are indicated. ***C.*** The *Cbln1* deletion in cerebellum significantly increases the amplitude of AMPAR-EPSCs at parallel-fiber synapses (left, sample traces of evoked AMPAR-EPSCs; middle, summary plot of AMPAR-EPSCs input-output curves; right, summary graph of the slope of AMPAR-EPSC input/output curves). ***D.*** The *Cbln1* deletion in cerebellum has no major effect on the coefficient of variation at parallel-fiber synapses, suggesting that it does not greatly change the release probability (left, sample traces of evoked AMPAR-EPSCs with 50 μA stimulus intensity; right, summary graph of the coefficient of variation of AMPAR-EPSCs). ***E*** *& **F***. Same as C & D, but recorded from Nrxn1^SS4+^ knockin mice in which ΔCre retains a constitutive expression of Nrxn1-SS4+ splice variants, whereas Cre converts the Nrxn1-SS4+ variants into constitutive Nrxn1-SS4-variants. ***G*** *& **H***. Same as E & F, but for Nrxn3^SS4+^ knockin mice in which ΔCre retains a constitutive expression of Nrxn1-SS4+ splice variants, whereas Cre converts the Nrxn1-SS4+ variants into constitutive Nrxn1-SS4-variants. Data are means ± SEM. Number of neurons/mice are indicated in bars. Statistical significance was assessed by two-way ANOVA or unpaired two-tailed t-test (*P ≤ 0.05, **P ≤ 0.01, and ***P ≤ 0.001).

These results appear to indicate that Nrxn3^SS4+^-Cbln1 complexes but not Nrxn1^SS4+^-Cbln1 complexes control parallel-fiber synapse properties in the cerebellum. Given the fact that both Nrxn1 and Nrxn3 are expressed in the cerebellum almost exclusively as SS4+ splice variants (Figure S2), this is surprising. To validate this conclusion, we again used the mouse lines of conditional genetic knockin of endogenous SS4+ and SS4-variants of Nrxn1 and Nrxn3. Measurements of parallel-fiber synaptic transmission revealed that the presynaptic Nrxn3^SS4+^ knockin fully phenocopies the Cbln1 cKO, whereas the Nrxn1^SS4+^ knockin had no effect (Figure 7E-7H). As before, none of these manipulations altered the coefficient of variation, suggesting that they did not influence the release probability (Figure 7F, 7H). These results confirm that the function of Cbln1 in cerebellum is dependent on presynaptic Nrxn3^SS4+^ signals and acts to control postsynaptic AMPAR responses at the PF-PC synapses (Figure 7).

## DISCUSSION

We previously showed that at CA1→subiculum synapses, signaling by Nrxn1^SS4+^ and Nrxn3^SS4+^ selectively enhances NMDAR-EPSCs and suppresses AMPAR-EPSCs, respectively, via a surprising common mechanism: Binding to Cbln2 that in turn binds to GluD1 (Dai et al., 2019 and 2021). The convergence of distinct Nrxn1^SS4+^ and Nrxn3^SS4+^ signals on the same Cbln2-GluD1 effectors to produce different downstream effects is unexpected, but was validated by the demonstration that distinct cytoplasmic GluD1 sequences transduced the differential Nrxn1^SS4+^ and Nrxn3^SS4+^ signals (Dai et al., 2021). These findings thus described a trans-synaptic signaling pathway regulating NMDARs and AMPARs, but raised new questions. In particular, given multiple lines of evidence suggesting a role for cerebellins in synapse formation (see Introduction) and given the fact that our experiments only manipulated mature neurons (Dai et al., 2019), the question arose whether Cbln2 may have additional functions in synapse formation at CA1→subiculum synapses during development, and whether additional roles of Cbln2 at CA1→subiculum synapses might have been redundantly occluded by the low levels of Cbln1 present. Most important, however, may be the question whether the Nrxn1^SS4+^-Cbln2 and Nrxn3^SS4+^-Cbln2 signaling pathways (and those of the closely related Cbln1 isoform) were specific to CA1→subiculum synapses, or whether they broadly operated in other synapses in brain as well. We have now addressed these questions. Our data suggest that at CA1→subiculum synapses, Cbln1 does not redundantly occlude a major additional function of Cbln2, that the Nrxn1^SS4+^-Cbln2 and Nrxn3^SS4+^-Cbln2 signaling pathways do not have additional synapse-formation functions, and that these signaling pathways are important regulators of NMDARs and AMPARs at multiple types of synapses in brain. Strikingly, however, these signaling pathways do not equally operate at all synapses, but are selectively present in subsets of synapses (Figure 8). The evidence supporting these conclusions can be summarized as follows.

**Figure 8:**
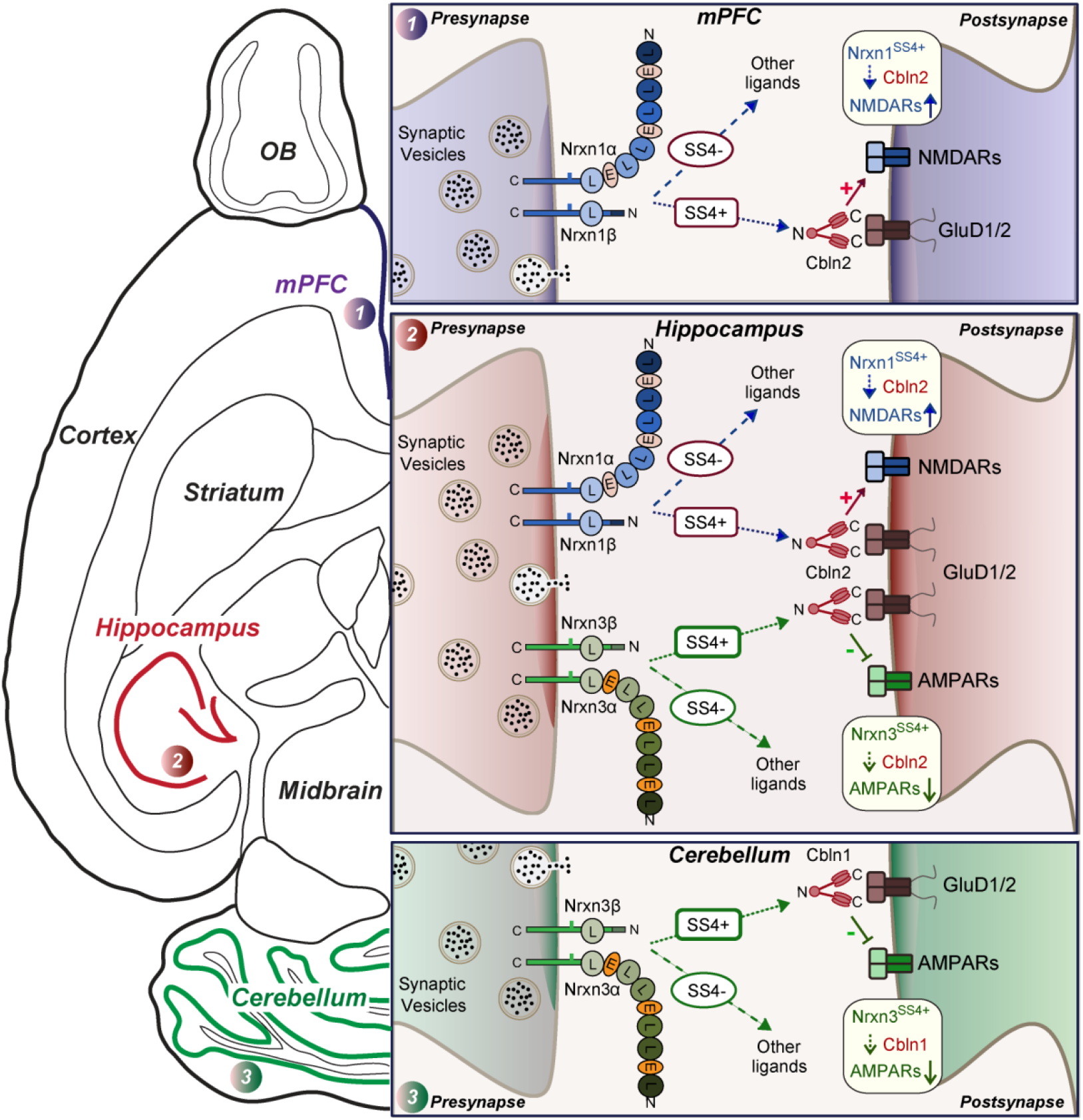
Schematic illustrating how Nrxn1^SS4+^-Cbln1/2 and Nrxn3^SS4+^-Cbln1/2 complexes control postsynaptic AMPARs and NMDARs in subicular, prefrontal, and cerebellar circuits. The schematic is based on data shown previously (Aoto et al., 2013; Dai et al., 2019 and 2021) and described here. Alternative splicing of presynaptic Nrxn1 and Nrxn3 at SS4 that controls their interactions with Cbln1/2 and thereby with postsynaptic GluD1/2 differentially regulates the postsynaptic content of AMPARs and NMDARs in different brain region. In the hippocampus, Nrxn1^SS4+^-Cbln1/2 complexes enhance NMDAR-EPSCs, whereas Nrxn3^SS4+^-Cbln1/2 complexes suppress AMPAR-EPSCs, with both types of complexes acting via GluD1/2. In the mPFC, Nrxn1^SS4+^-Cbln1/2 complexes also enhance NMDAR-EPSCs, but Nrxn3^SS4+^-Cbln1/2 complexes have no effect. In the cerebellum, conversely, Nrxn3^SS4+^-Cbln1/2 complexes suppress AMPAR-EPSCs, whereas now Nrxn1^SS4+^-Cbln1/2 complexes have no effect. These results indicate that Nrxn1^SS4+^-Cbln1/2 and Nrxn3^SS4+^-Cbln1/2 complexes perform universal functions in regulating AMPARs and NMDARs, respectively, but that these regulatory signaling pathways are differentially expressed in different types of synapses.

First, we showed that a constitutive deletion of *Cbln2* operating throughout development has the same effect as the conditional post-developmental deletion of *Cbln2* (Figure 1-3). Both produced the same enhancement of AMPAR-EPSCs (up to 100% increase) and the same suppression of NMDAR-EPSCs (up to 40% decrease) without a change in synapse numbers. Consistent with its broad effect on synapses, the *Cbln2* deletion also severely impaired contextual learning (Figure 2). Moreover, the constitutive deletion of *Cbln2* occluded the dominant effects of Nrxn1^SS4+^ and Nrxn3^SS4+^ signaling on NMDARs and AMPARs, respectively (Figure 4), confirming that Nrxn1^SS4+^ and Nrxn3^SS4+^ operate upstream of Cbln2.

Second, we examined whether the function of Cbln2 is the same in the two types of CA1→subiculum synapses that are formed on burst- and regular-spiking neurons and that exhibit quite distinct properties (Wojtowicz et al., 2010; Wozny et al., 2008a and 2008b). In both synapse types, the constitutive and the conditional *Cbln2* deletion caused the same increase in AMPAR-EPSCs and the same decrease in NMDAR-EPSCs (Figure 1, S1). The two types of subiculum synapses differ in their form of LTP (Wojtowicz et al., 2010; Wozny et al., 2008a and 2008b). Notably, the *Cbln2* deletion blocked the NMDAR-dependent LTP of synapses on regular-spiking neurons, possibly via an induction impairment, without affecting the cAMP-dependent LTP in burst-spiking neurons (Figure 1). Since the former type of LTP is postsynaptic and latter presynaptic, these findings agree with the conclusion of a postsynaptic regulatory effect of Cbln2 signaling.

Third, we investigated the possibility that low-level expression of Cbln1 in the subiculum might redundantly compensate for Cbln2 in an additional function besides the regulation of AMPARs and NMDARs, causing such a function to become occluded in the *Cbln2* KO mice. To explore this possibility, we analyzed *Cbln1/2* double KO mice, but identified substantially the same phenotype as in *Cbln2* single KO mice (Figure 5). Thus, it seems unlikely that low-level expression of Cbln1 prevents manifestation of an additional Cbln2 function.

Fourth, we tested the possible function of the Nrxn1^SS4+^-Cbln2 and Nrxn3^SS4+^-Cbln2 signaling pathways at L2/3→L5/6 synapses in the mPFC, focusing on *Cbln2* because it is expressed at higher levels than *Cbln1* in the mPFC (Figure S3). We observed that the *Cbln2* deletion caused a suppression of NMDAR-EPSCs, but did not enhance AMPAR-EPSCs (Figure 6). This observation suggests that only the Nrxn1^SS4+^-Cbln2 but not the Nrxn3^SS4+^-Cbln2 signaling pathway operates in the mPFC synapses. Consistent with this conclusion, we found that the Nrxn1^SS4+^ switch to Nrxn1^SS4-^ selectively downregulated NMDARs because only Nrxn1^SS4+^ but not Nrxn1^SS4-^ can bind to Cbln2, whereas different from CA1→subiculum synapses, the Nrxn3^SS4+^ switch to Nrxn3^SS4-^ had no effect on AMPARs (Figure 6). A recent study discovered a hominin-specific increase in *Cbln2* expression in the prefrontal cortex in primates (Shibata et al., 2021). The study suggested that this finding might imply a higher synapse density in hominins, but our data suggest that this interesting observation is associated with an increased NMDAR expression at synapses in hominid evolution.

Fifth and finally, we examined parallel-fiber synapses in the cerebellum, the synapses at which Cbln1 was discovered and which do not express functional NMDARs (Llano et al., 1991; Perkel et al., 1990). Cbln1 has a well-characterized function at these synapses in maintaining synapse stability and enabling long-term synaptic plasticity (Kashiwabuchi et al., 1995; Kurihara et al., 1997; Hirai et al., 2005; Takeuchi et al., 2005). Strikingly, we found that the post-developmental conditional deletion of *Cbln1* at these synapses also significantly increased AMPAR-EPSCs, and that the induced switch from Nrxn3^SS4+^ to Nrxn3^SS4-^ had the same effect on AMPAR-EPSCs, whereas the Nrxn1^SS4+^ switch to Nrxn1^SS4-^ had no effect (Figure 7). These experiments suggest that in parallel-fiber synapses of the cerebellum, Nrxn3^SS4+^-Cbln1 signaling controls AMPARs similar to the action of Nrxn3^SS4+^-Cbln2 signaling at CA1→subiculum synapses (Dai et al., 2019). At first glance, these results seem to contradict previous studies on the deletions of *Cbln1* and its GluD2 receptor in cerebellum, which cause a loss of parallel-fiber synapses (reviewed in Yuzaki, 2018; Yuzaki and Aricescu, 2017). However, this loss affects less than half of all synapses, while the remaining synapses are abnormal since they can’t undergo LTD. More importantly, this loss only occurs after an initially apparently normal formation of synapses (Kurihara et al., 1997), and the GluD2 deletion was also previously shown to induce an increase in AMPARs at parallel-fiber synapses (Yamasaki et al., 2011), replicating our observations with the *Cbln1* deletion since *Cbln1* is the major binding partner to GluD2 (Figure 7).

Figure 8 illustrates the richness of regulatory mechanisms that control the postsynaptic levels of AMPARs and NMDARs via presynaptic expression of neurexins and cerebellins. In our studies, the changes in synaptic transmission induced by disrupting neurexin-cerebellin signaling are large, resulting in major alterations in the information processing of any circuit containing affected synapses. Since both cerebellin expression (Hrvatin et al., 2018; Ibata et al., 2019) and neurexin alternative splicing at SS4 (Iijima et al., 2011; Ding et al., 2017; Flaherty et al., 2019) may be activity-dependent, the unexpected signaling mechanism we describe likely also mediates activity-dependent plasticity. Thus, activity-dependent gene expression changes in a pre- or postsynaptic neuron may regulate the AMPAR- and NMDAR-composition via Nrxn1^SS4+^/Nrxn3^SS4+^→Cbln signaling. This type of AMPAR and NMDAR plasticity, which has not been previously identified, suggests a novel mechanism of circuit plasticity that may contribute to fundamental brain functions such as learning and memory (Silver et al., 2010; Josselyn and Tonegawa, 2020).

Needless to say, our findings raise major new questions. The current data at best are the beginning of an understanding of how neurexin-cerebellin signaling shapes synapses. Among major questions, most prominent may be the puzzle of why Nrxn3^SS4+^ has no effect on mPFC synapses. It is expressed in the mPFC as the SS4+ the variant but doesn’t regulate AMPARs, suggesting it has a different function that is independent of Cbln2. In contrast, it is easier to understand why Nrxn1^SS4+^ doesn’t regulate NMDARs at parallel-fiber synapses since these synapses lack functional NMDARs (Llano et al., 1991; Perkel et al., 1990), but this finding also raises the question whether Nrxn1^SS4+^ has another currently unknown function at these synapses. Neurexins can likely operate at the same synapses via binding to different ligands (Wang et al., 2021), a fascinating amplification of their functions that may also apply to parallel-fiber synapses. A further question is how the function of Cbln1 and Nrxn3^SS4+^ in regulating AMPARs relates to the well-described parallel-fiber synapse loss in constitutive Cbln1 KO mice. It is possible that an overactivation of AMPARs leads to synaptotoxicity that destroys parallel-fiber synapses; another plausible explanation could be that at cerebellar parallel-fiber synapses, Cbln1 has additional functions that are not operative for cerebellins in subiculum synapses. Future studies will have to explore these intriguing questions.

In summary, our data spanning diverse genetic manipulations in multiple brain regions establish a general function for *Cbln1* and *Cbln2* in the trans-synaptic regulation of NMDARs and AMPARs by presynaptic Nrxn1^SS4+^ and Nrxn3^SS4+^, respectively. Remarkably, this signaling pathway differentially operates in different neural circuits, creating a panoply of synaptic regulatory mechanisms that are inherently plastic and enhance the activity-dependent capacity for information processing by neural circuits.

## METHODS

### Mice

The Cbln1 conditional KO and Cbln2 conditional/constitutive KO mouse lines were described in Seigneur and Südhof (2017). SS4+ conditional knockin (cKI) mice of *Nrxn1* and *Nrxn3* were described previously (Aoto et al., 2013; Dai et al., 2019; Dai et al., 2021). All mice above were maintained on a mixed C57BL/6/SV129/CD1 (wild type) background. Primers (IDT) are used for genotyping are as follows: Nrxn1-SS4+, forward: 5’- AGACAGACCCGAACAACCAA-3’, reverse: 5’-TGCTAGGCCTATTTCAGATGCT-3’; Nrxn3- SS4+, forward: 5’-CTCCAACCTGTCATTCAAGGG-3’, reverse: 5’- CTACGGGCCGGTTATATTTG-3’; Cbln1, LoxP forward: 5’-TAGGG TGGACAGAGAAAAGG-‘3, LoxP reverse: 5’- CTTCTAATCTGTCCTGACCACA-‘3; Cbln2, LoxP forward: 5’-TAAAAGACAGTCCAGAGTTTTAGTC-3’, LoxP reverse: 5’- TCAAATAGAGAGGAGTAAGCACA-3’, and Recombined reverse: 5’- TTTCCTTGAAGGACTCCAATAG-3’. All mouse studies were performed according to protocols approved by the Stanford University Administrative Panel on Laboratory Animal Care. In all studies, we examined littermate male or female mice.

### Single-molecule RNA FISH

As described in our previous study (Dai et al., 2021), P30 Wild type BL6 mice were euthanized with isofluorane and followed by transcardial perfusion with ice cold PBS. The brain were quickly dissected and embedded in OCT (Optimal Cutting Temperature) solution on dry ice. Horizontal sections with 16 um thickness were cut by using Leica CM3050-S cryostat, mounted directly onto Superfrost Plus slides and stored in -80 °C until use. Single-molecule FISH for Cbln1 (Cat# 538491-C2) and Cbln2 (Cat# 428551) mRNA was performed using the multiplex RNAscope platform (Advanced Cell Diagnostics) according to manufacturer instructions. Fluorescent microscopy images were acquired at 20x magnification using Olympus VS120 slide scanner.

### Semi-quantitative RT-PCR

For semi-quantitative RT-PCR measurements of neurexin SS4 alternative splicing (Liakath-Ali and Sudhof, 2021), total RNA was extracted using TRIzol and cDNA was synthesized using the SuperScript III First-Strand Synthesis System (Invitrogen) according to the manufacturer’s instructions. PCR primers to detect Nrxn-SS4 isoforms (Forward, reverse): Nrxn1SS4, CTGGCCAGTTATCGAACGCT, GCGATGTTGGCATCGTTCTC; Nrxn2SS4, CAACGAGAGGTACCCGGC, TACTAGCCGTAGGTGGCCTT; Nrxn3SS4, ACACTTCAGGTGGACAACTG, AGTTGACCTTGGAAGAGACG; β-actin, TTGTTACCAACTGGGACGACA, TCGAAGTCTAGAGCAACATAGC.

### DNA constructs and Viruses

hSyn-Cre-eGFP, hSyn-ΔCre-eGFP, CAG-Cre-eGFP, CAG-ΔCre-eGFP, full-length Nrxn1β^SS4+^, Nrxn1β^SS4-^, Nrxn3β^SS4+^, and Nrxn3β^SS4-^ were cloned into AAV-DJ vector (Xu et al., 2012; Aoto et al., 2013; Dai et al., 2019) for in vivo Cre-recombination or overexpression as previously described (Dai et al., 2019). The overexpression levels mediated by the viruses were quantified in microdissected brain tissue (please see details in Dai et al., 2019).

### Slice Electrophysiology

As previously described, electrophysiological recordings from acute hippocampal slices (Dai et al., 2019; Dai et al., 2021) or prefrontal cortex (Xu et al., 2012) or cerebellum (Zhang et al., 2015) were essentially performed. In brief, slices were prepared from Cbln2^+/+^ and Cbln2^-/-^ mice at P35-42, or from all other mice at 2-3 weeks after stereotactic infection of AAVs (encode Cre, ΔCre, and various β-neurexins). Horizontal hippocampal slices (300 μm thickness) and Coronal prefrontal cortex (PFC) slices (250 μm thickness) were cut in a high sucrose cutting solution containing (in mM) 85 NaCl, 75 sucrose, 2.5 KCl, 1.3 NaH_2_PO_4_, 24 NaHCO_3_, 0.5 CaCl_2_, 4 MgCl_2_ and 25 D-glucose. Sagittal cerebellum slices were sectioned in a low calcium solution containing (in mM) 125 mM NaCl, 2.5 mM KCl, 3 mM MgCl_2_, 0.1 CaCl_2_, 1.25 NaH_2_PO_4_, 25 NaHCO_3_, 3 mM myo-inositol, 2 mM Na-pyruvate, 0.4 mM ascorbic acid, and 25 D-glucose. Slices were equilibrated in ACSF at 31 °C for 30 min, followed by room temperature for an hour. Hippocampal or PFC Slices were then transferred to a recording chamber containing ACSF solution maintained at 30.5°C (in mM): 120 NaCl, 2.5 KCl, 1 NaH_2_PO_4_, 26.2 NaHCO_3_, 2.5 CaCl_2_, 1.3 MgSO_4_-7 H_2_O, 11 D-Glucose, ∼290 mOsm. Cerebellum slices were then transferred to a recording chamber containing ACSF solution maintained at 30.5°C (in mM): 125 mM NaCl, 2.5 mM KCl, 1 mM MgCl_2_, 2 CaCl_2_, 1.25 NaH_2_PO_4_, 25 NaHCO_3_, 3 mM myo-inositol, 2 mM Na-pyruvate, 0.4 mM ascorbic acid, and 25 D-glucose. To induce evoked synaptic responses in subiculum, a nichrome stimulating electrode was placed at the most distal portion of hippocampal CA1 region as shown in our previous studies (Dai et al., 2019; Dai et al., 2021). The firing type of subiculum neurons (burst-spiking vs. regular-spiking) was identified by injecting a depolarizing current immediately after breaking in and monitoring action potential patterns in current-clamp mode as previously described (Aoto et al., 2013; Dai et al., 2019). To induce evoked synaptic responses in mPFC, the electrode was placed at the border of L1 and L2/3 layer as illustrated in Figure 6B and the L5/6 layer pyramidal neurons were recorded (Fenelon et al., 2011). To induce evoked synaptic responses in cerebellum, the electrode was placed at the parallel fibers in the distal molecular layer as illustrated in Figure 7B and the purkinje neurons were recorded (Zhang et al., 2015). AMPAR-EPSCs input/output curves, AMPAR/NMDAR ratios, NMDAR input/output curves, LTP, and mEPSCs (holding potentials = -70 mV for AMPAR-EPSCs, +40 mV for NMDAR-EPSCs, and +60 mV for NMDAR mEPSCs) were recorded with an internal solution containing (in mM): 117 Cs-methanesulfonate, 15 CsCl, 8 NaCl, 10 TEA-Cl, 0.2 EGTA, 4 Na2-ATP, 0.3 Na2-GTP, 10 HEPES, pH 7.3 with CsOH (∼300 mOsm). All recordings were performed in the presence of 50 μM picrotoxin for AMPAR-EPSCs, 50 μM picrotoxin and 10 μM CNQX for NMDAR-EPSCs, and 50 μM picrotoxin and 0.5 μM TTX for mEPSCs. Paired-pulse ratios were monitored with interstimulus intervals of 20–2000 ms. LTP was induced by four tetani of 100 Hz stimulus trains applied for 1 s with 10 s intervals under voltage-clamp mode (holding potential = 0 mV). Pre-LTP (averaging last 5 mins as baseline) and post-LTP (averaging the last 5 mins) were recorded at 0.1 Hz. Paired-pulse ratios were measured with 40 ms interstimulus intervals before and after LTP. Measurements of the AMPAR/NMDAR ratios were performed in 50 μM picrotoxin at holding potentials of -70 mV (AMPAR-EPSCs) or +40 mV (NMDAR-EPSCs, quantified at 50 ms after the stimulus). All slopes of input/output ratio were calculated from 10-50 μA of input current except the cerebellum that was calculated from 10-100 μA of input current. All data were analyzed with the Igor software (WaveMetrics). Miniature events were handpicked with a threshold of 5 pA by using the Igor software (Dai et al., 2015).

### Stereotactic Injections

Stereotactic injections of AAV or Lentivirus into mice at P21 were performed essentially as described (Xu et al., 2012; Aoto et al., 2013; Dai et al., 2019; Dai et al., 2021). Briefly, P21 mice were anesthetized with Avertin, and viruses were injected using a stereotactic instrument (David Kopf) and a syringe pump (Harvard Apparatus) with ∼0.85 μl of concentrated virus solution (10^8-9^ TU) at a slow rate (0.1ul/min) into the CA1 region of the intermediate hippocampus (Bregma coordinates (mm): AP: −3.1, ML: ± 3.4, DV: −2.5) or with ∼0.4 μl of virus into subiculum region (Bregma coordinates (mm): AP: −3.3, ML: ± 3.3, DV: −2.5) or with ∼0.8 μl of virus into mPFC region (Bregma coordinates (mm): AP: +1.25, ML: ± 0.3, DV: −1.0 mm and -1.5 mm received both 0.4 μl of virus) or with ∼0.8 μl of virus into cerebellum lobe4-5 region (Bregma coordinates (mm): AP: -6.35, ML: ± 0.5, DV: −1.5 mm received both 0.4 μl of virus. After infection, viral mediated expression was confirmed by the presence of eGFP. Images (Figure 4F, 6B, and 7B) were taken using a Nikon confocal microscope (A1Rsi) with a 10x objective (PlanApo, NA1.4) with 1024×1024 pixel resolution. The fluorescence of all slices prepared for physiology was confirmed under a fluorescence microscope (Olympus).

### Immunohistochemistry

For hippocampal cryosections were performed as described (Dai et al., 2019; Dai et al., 2021). Briefly, mice were anesthetized with isoflurane and perfused with 10 ml PBS followed by 30 ml 4% PFA in 1x PBS using a perfusion pump (2 ml/min). Whole brains were dissected out and kept in PFA for 6 hours, then post-fixed in 30% sucrose (in 1× PBS) for 24 h-48 h at 4°C. Horizontal brain sections (30 μm) were collected at -20°C with a cryostat (Leica CM1050). Sections were washed with PBS and incubated in blocking buffer (0.3 % Triton X-100 and 5% goat serum in PBS) for 1 h at RT, and incubated overnight at 4 °C with primary antibodies diluted in blocking buffer (anti-vGluT1, 1:1000, guinea pig, Millipore and anti-MAP2, 1:1000, rabbit, Millipore). Sections were washed 3 times for 10 mins each in 1x PBS, followed by treatment with secondary antibodies (1:1000, Alexa 405, Alexa 647) at 4°C overnight, then washed 3 times for 10 mins each with 1x PBS. All incubations were performed with agitation. All sections were then mounted on superfrost slides and covered with Fluoromount-G as previously described. Serial confocal z-stack images (1 μm step for 10 μm at 1024 x 1024 pixel resolution) were acquired using a Nikon confocal microscope (A1Rsi) with a 60x oil objective (PlanApo, NA1.4). All acquisition parameters were kept constant among different conditions within experiments. For data analysis (n≥3 animals per condition), maximum intensity projections were generated for each image, and average vGlut1 intensity (mean ± S.E.M) calculated from the entire area of subiculum (object size range 0.05-0.21 mm^2^). An example cerebellum slice was stained with vGluT1 (anti-vGluT1, 1:1000, guinea pig, Millipore) and Calbindin (anti-calbindin, 1:2000, mouse, Sigma).

### Immunoblotting

Immunoblotting was performed as described previously (Seigneur and Sudhof, 2018; Patzke et al., 2019; Dai et al., 2021; Patzke et al., 2021). Briefly, dissected hippocampal tissue were homogenized in Laemmli buffer (12.5 mM Tris-HCl, pH 6.8, 5 mM EDTA, pH 6.8, 143 mM β-mercaptoethanol, 1% SDS, 0.01% bromophenol blue, 10% glycerol), boiled and separated by SDS–PAGE at 100 V for about 1.3 h, then transferred onto nitrocellulose membranes using the Trans-Blot Turbo transfer system (Bio-Rad). Membranes were then blocked with 5% milk in TBS containing 0.1% Tween 20 (TBST) at RT for 1h, and then incubated in primary antibody overnight at 4°C. Membranes were washed 3X with TBST, then incubated in fluorescent labeled secondary antibodies (donkey anti-rabbit IR dye 680/800CW, 1:10000; donkey anti-mouse IR dye 680/800CW, 1:10000; and donkey anti-guinea pig IR dye 680RD, 1:10000; LI-COR Bioscience). Membranes were scanned using an Odyssey Infrared Imager and analyzed with the Odyssey software (LI-COR Biosciences). Intensity values for each protein were first normalized to actin then to the control sample. The antibodies used are as follows: anti-Neuroligin-1 mouse (1:500; Südhof lab; 4F9), anti-β-actin mouse (1:10000; Sigma-Aldrich; Cat# A1978), anti-PSD95 rabbit (1:500; Südhof lab; L667), anti-Synapsin rabbit (1:1000; Südhof lab; E028), anti-CASK mouse (1:1000; BD Transduction Laboratories; Cat# 610782), anti-Neurexin rabbit (1:500; Südhof lab; G394), anti-GAD65 mouse (1:500; DSHB; Cat# mGAD6-a), anti-Synaptotagmin-1 mouse (1:1000, Südhof lab; CL41.1), and anti-vGluT1 guinea pig (1:1000; Millipore; Cat# AB5905).

### Two-chamber avoidance test

Littermate Cbln2 WT and Cbln2 KO male mice were generated from crossing heterozygous Cbln2^+/-^ mice. Mice were handled daily for 5 days prior to behavioral experiments starting at P45. Mice were maintained with a normal 12/12 hr daylight cycle, and analyzed in the assay sequence and at the time shown in figure 2A. The modified protocol was performed as described previously (Dai et al., 2019) and was based on previous studies (Ambrogi Lorenzini et al., 1984; Cimadevilla et al., 2001; Qiao et al., 2014). Briefly, two chambers (left and right) were designed with different visual cues (figure supplement 4C) under dim light with a gate between them. The right chamber has a foot shock with electric current (intensity: 0.15 mA, duration: 2s). Mice can explore both chambers freely. At the training day, mice will be put in left chamber. Once they go to the right chamber, they will get a foot shock after a 2 s delay. In this case, they will return back immediately to the left chamber. This is one trial of learning. It may come as another trial, once they visit right chamber again. This training process will be completed until mice are able to stay in left “safe” chamber more than 2 mins. After 1 day and 7 days, they will be tested by putting back into left chamber to record latency to enter the right chamber and the number of entries in 2 mins. Using this approach, two groups of Cbln2 WT and KO mice were tested. All behavior assays were carried out and analyzed by researchers blindly.

### Quantification and statistical analysis

All data are shown as means ± SEMs, with statistical significance (* = *p*<0.05, ** = *p*<0.01 and = *p*<0.001) determined by Student’s t-test or two-way analysis of variance (ANOVA). Non-significant results (*p*>0.05) are not specifically identified.

## ACKNOWLEDGEMENTS

This study was supported by a grant from the NIMH (MH052804 to T.C.S.) and fellowships to K.L-A. from the European Molecular Biology Organization (ALTF 803-2017) and the Larry L. Hillblom Foundation (2020-A-016-FEL).

## AUTHOR CONTRIBUTIONS

J.D. performed all experiments and analyzed all data except for the in situ hybridization experiment and semi-quantitative RT-PCR of alternative splicing variants that were performed by K.L-A., and S.G. contributed mouse behavioral studies and genotyping experiments. J.D. and T.C.S. conceptualized the project, designed the experiments and wrote the manuscript with inputs from all authors.

## MATERIALS AVAILABILITY

All reagents produced in this study, including recombinant DNA plasmids and mouse lines, are openly distributed to the scientific community and freely shared upon request.

## DATA AVAILABILITY

All numerical data and *P* values within this study have been included in the source data.

## KEY RESOURCES TABLE

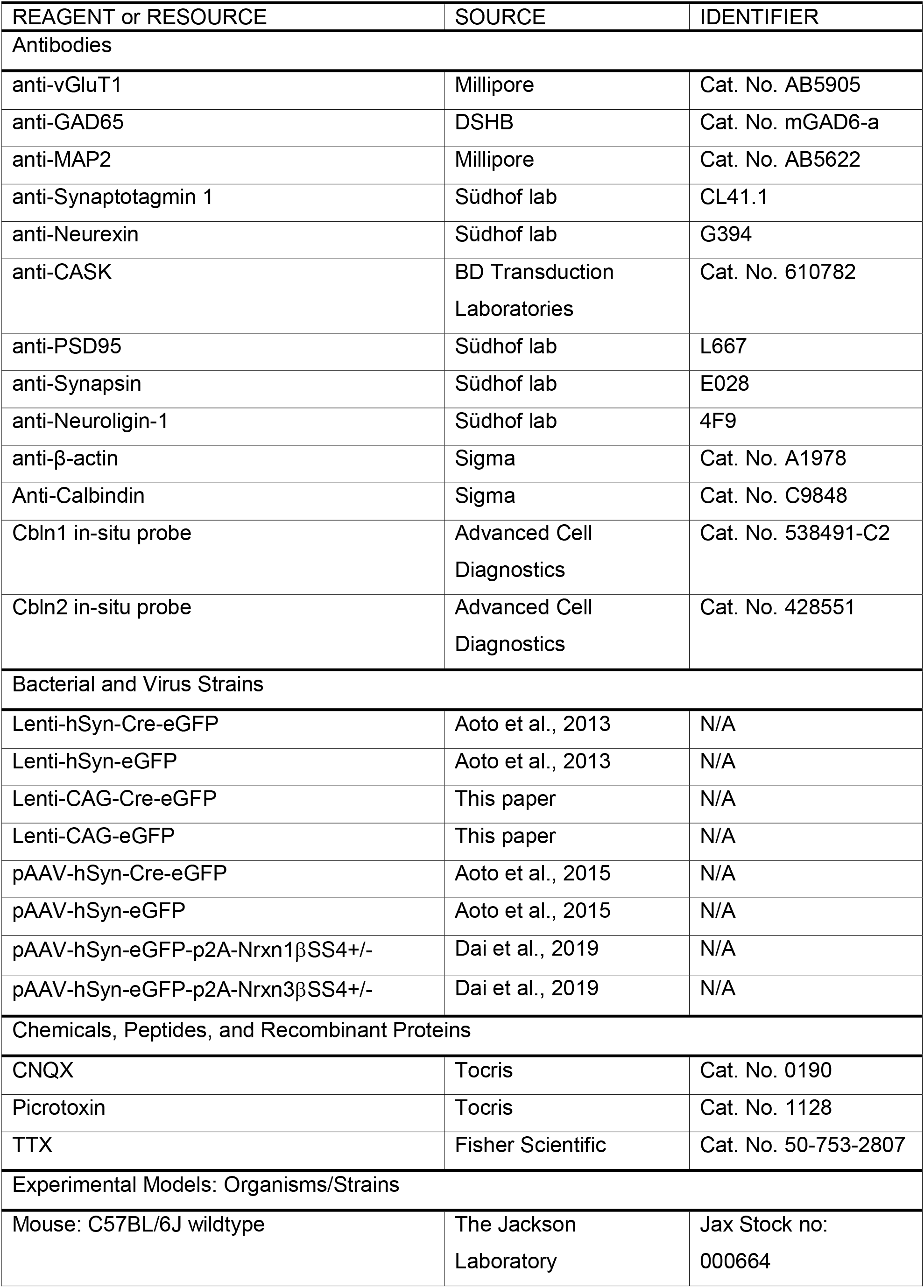
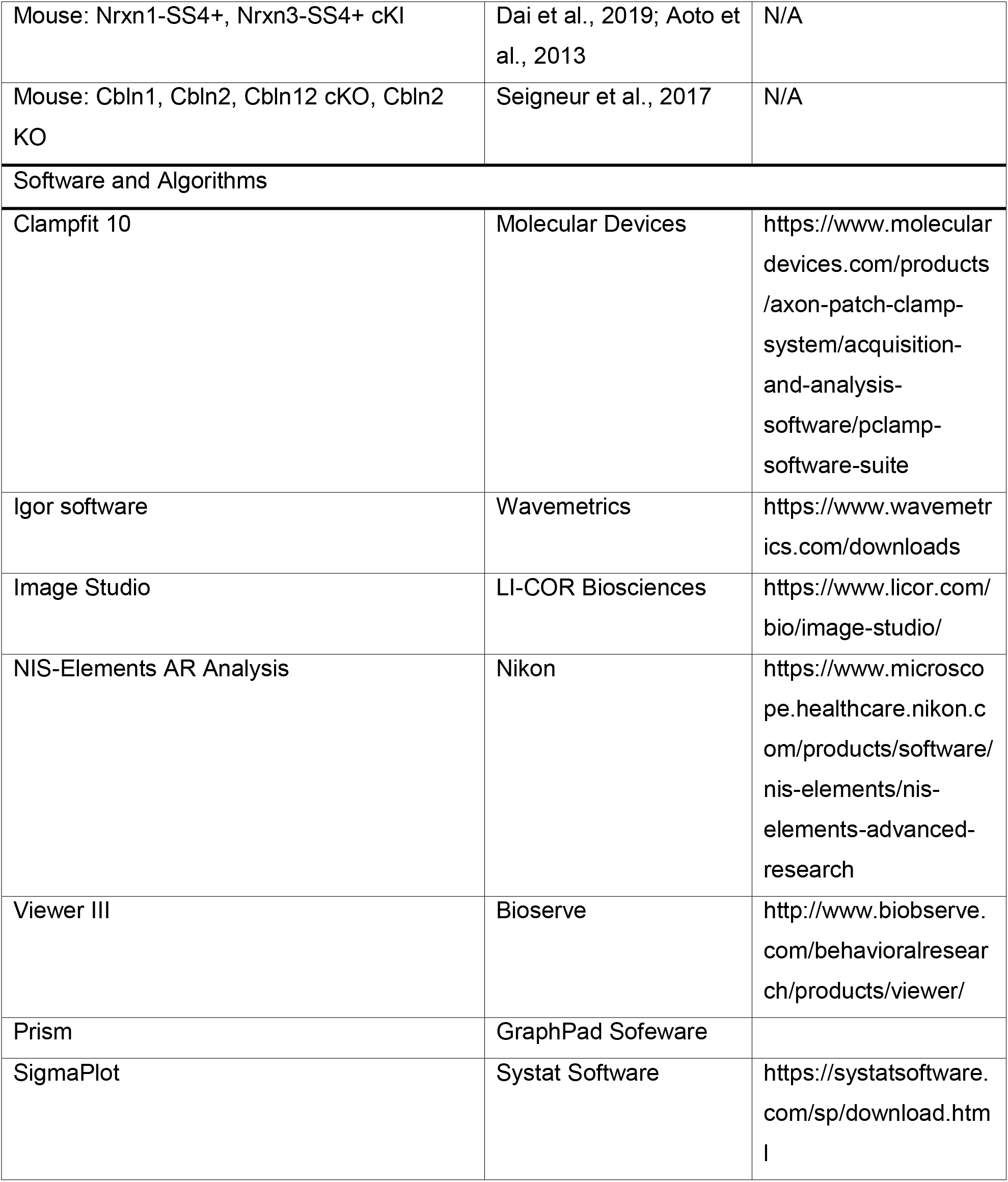

## SUPPLEMENTAL FIGURES and FIGURE LEGENDS

**Figure S1:**
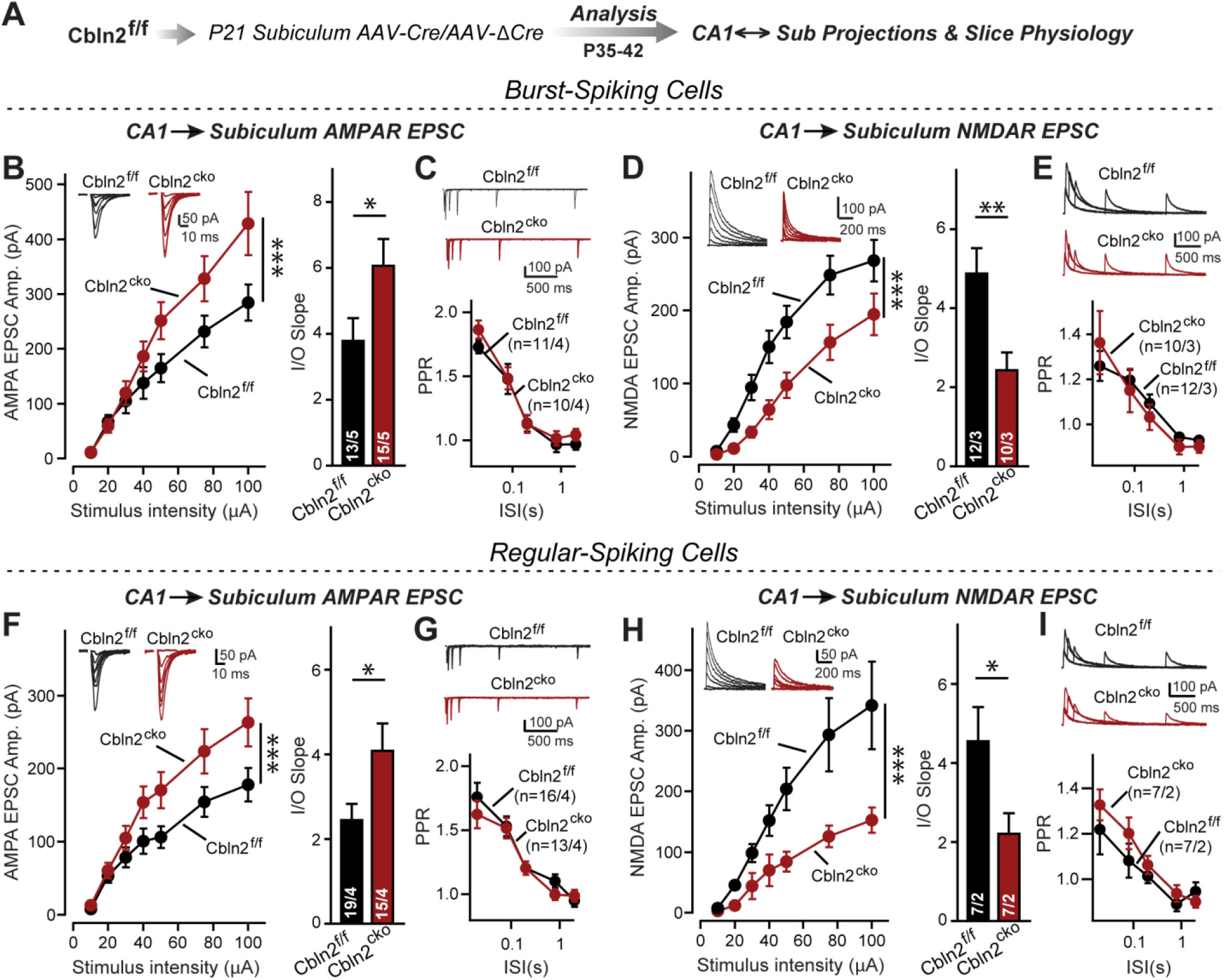
Conditional *Cbln2* KO in the subiculum produces the same phenotype as the constitutive *Cbln2* KO at the two different types of CA1→subiculum synapses that are formed on burst- and regular-spiking neurons. ***A.*** Experimental design for the generation and analysis of littermate control and conditional *Cbln2* KO mice. AAVs encoding Cre or ΔCre (as a control) were stereotactically injected into the subiculum of conditional *Cbln2* KO mice at P21, and mice were analyzed by slice physiology 2-3 weeks later. ***B****-**E***. Input/output measurements of evoked AMPAR-EPSCs (B) and NMDAR-EPSCs (D) recorded from burst-spiking neurons in acute subiculum slices reveal that the conditional *Cbln2* KO enhances AMPAR-EPSCs (B) without changing the paired-pulse ratio of AMPAR-EPSCs (C) but suppresses NMDAR-EPSCs (D), again without changing the paired-pulse ratio of NMDAR-EPSCs (E), in burst-spiking neurons. Sample traces are shown above the respective summary plots and graphs. ***F****-**I.*** Same as B-E, but recorded from regular-spiking neurons. Note that the conditional *Cbln2* KO phenotype is identical between burst- and regular-spiking neurons. Data are means ± SEM. Number of neurons/mice are indicated in bars. Statistical significance was assessed by unpaired two-tailed t-test or two-way ANOVA (*P≤0.05, **P≤0.01, and ***P≤0.001).

**Figure S2:**
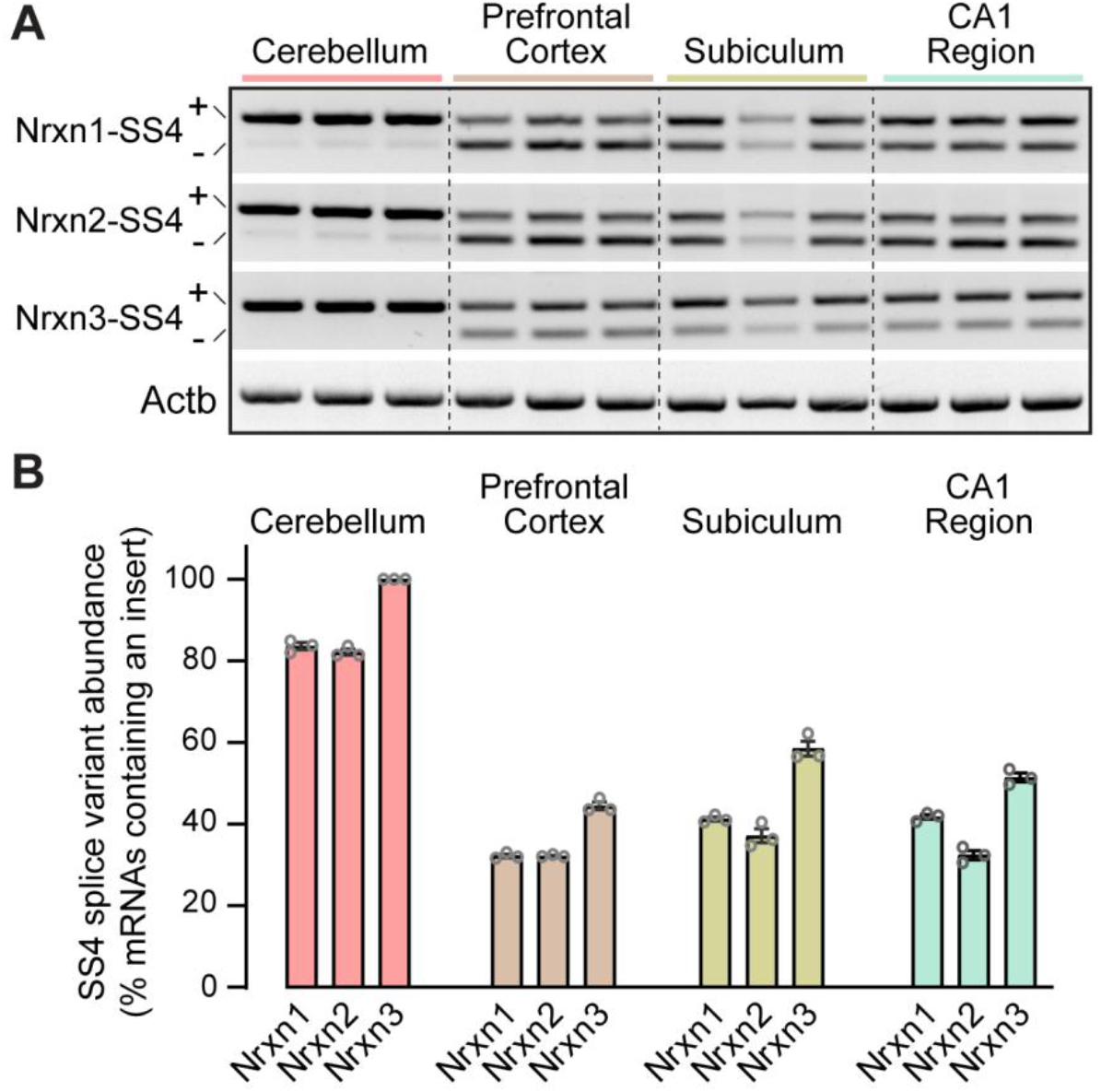
Analysis of neurexin SS4 alternative splicing reveals that whereas all neurexins are almost exclusively expressed as SS4+ variants in the cerebellum, in other brain regions a mixture of SS4+ and SS4-variants is observed. ***A.*** Sample gels of amplified DNA obtained by RT-PCR of mRNAs from the indicated brain regions. RT-PCR was carried out with primers flanking SS4; as a result, in most brain regions two bands are observed that correspond to mRNAs containing (upper bands) and lacking SS4 (lower bands). Please also see original full-sized gels in Figure supplement 2-source data 1. ***B.*** Quantification of the prevalence of SS4+ variants of the three neurexins in the indicated brain regions. Data are means ± SEM (n = 3).

**Figure S3:**
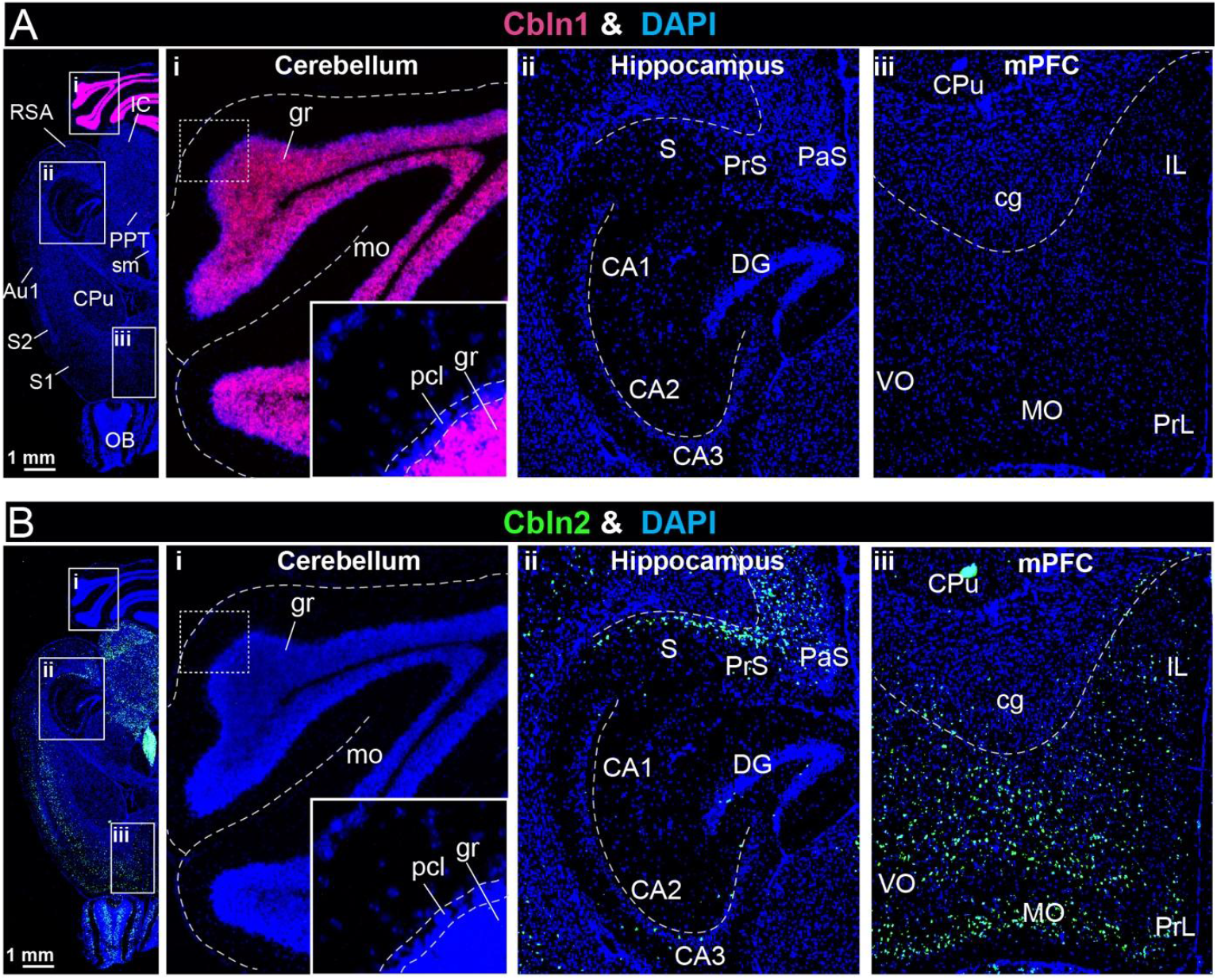
Analyses of the region-specific expression patterns of Cbln1 and Cbln2 using single-molecule RNA in situ hybridization (A & B) ***A*** *& **B***. Single-molecule in-situ hybridization analysis of Cbln1 (A) and Cbln2 mRNAs (B) reveals highly restricted expression patterns in brain (left, overview of horizontal mouse brain sections hybridized for Cbln1 or Cbln2 mRNAs; right, representative images for Cbln1 and Cbln2 in the cerebellum (i), hippocampal formation (ii), and mPFC (iii)). Note that Cbln1 is highly expressed only in the cerebellum, whereas Cbln2 is most abundant in the subiculum and mPFC (abbreviations used: RSA, retrosplenial agranular cortex; IC, inferior colliculus; PPT, posterior pretectal nucleus; sm, stria medullaris; Au1, primary autitory area; S1 & S2, primary and secondary somatosensory cortex; OB, olfactory bulb; gr, granular layer; mo, molecular layer; pcl, purkinje cell layer; S, subiculum; PrS, presubiculum; Pas, parasubiculum; CA1, 2, 3, cornu ammonis 1, 2, 3; DG, dentate gyrus; CPu, caudate putamen (striatum); cg, cingulum; IL, infralimbic cortex; VO, ventro orbital cortex; MO, medial orbital cortex; PrL, prelimbic cortex).

